# Neuronal non-CG methylation is an essential target for MeCP2 function

**DOI:** 10.1101/2020.07.02.184614

**Authors:** Rebekah Tillotson, Justyna Cholewa-Waclaw, Kashyap Chhatbar, John Connelly, Sophie A. Kirschner, Shaun Webb, Martha V. Koerner, Jim Selfridge, David Kelly, Dina De Sousa, Kyla Brown, Matthew J. Lyst, Skirmantas Kriaucionis, Adrian Bird

**Author notes:** These authors share first authorship on this work (Rebekah Tillotson, Justyna Cholewa-Waclaw). Genetics and Genome Biology Program, The Hospital for Sick Children, The Peter Gilgan Centre for Research and Learning, Toronto, ON M5G 0A4, Canada.

## Abstract

DNA methylation is implicated in neuronal biology via the protein MeCP2, mutation of which causes Rett syndrome. MeCP2 recruits the NCOR1/2 corepressor complexes to methylated cytosine in the CG dinucleotide, but also to non-CG methylation, which is abundant specifically in neuronal genomes. To test the biological significance of its dual binding specificity, we replaced the MeCP2 DNA binding domain with an orthologous domain whose specificity is restricted to mCG motifs. Knock-in mice expressing the domain-swap protein displayed severe Rett syndrome-like phenotypes, demonstrating that interaction with sites of non-CG methylation, specifically the mCAC trinucleotide, is critical for normal brain function. The results support the notion that the delayed onset of Rett syndrome is due to the late accumulation of both mCAC and its reader MeCP2. Intriguingly, genes dysregulated in both *Mecp2*-null and domain-swap mice are implicated in other neurological disorders, potentially highlighting targets of particular relevance to the Rett syndrome phenotype.

## INTRODUCTION

Heterozygous loss-of-function mutations in the X-linked *MECP2* gene result in Rett syndrome (RTT), a neurological disorder affecting approximately 1 in 10,000 live female births (Amir et al., 1999). MeCP2 protein was initially identified by its ability to bind methylated CG dinucleotides and shown to repress transcription via recruitment of histone deacetylase complexes (Lewis et al., 1992; Nan et al., 1997, 1998). Recent evidence indicates that MeCP2 negatively regulates the expression of hundreds of genes via the recruitment of the HDAC3-containing NCOR1/2 corepressor complexes (Cholewa-Waclaw et al., 2019; Gabel et al., 2015; Lagger et al., 2017; Lyst et al., 2013). In the absence of functional MeCP2, indirect mechanisms also lead to the downregulation of hundreds more genes, perhaps connected with a global reduction in total RNA levels (Lagger et al., 2017; Li et al., 2013; Yazdani et al., 2012). This multitude of subtle changes to neuronal gene expression is thought to underlie Rett syndrome.

In addition to canonical mCG dinucleotides in duplex DNA, MeCP2 targets 5-methylcytosine in a non-CG context (or mCH, where H = A, C or T). This study focuses on the biological significance of this dual DNA binding specificity. During early mammalian embryogenesis, methylation of CG dinucleotides is at first depleted, then re-established, reaching high levels in the bulk genome while absent at unmethylated CpG islands (Bird et al., 1985; Deaton and Bird, 2011). *De novo* CG methylation is laid down by the DNA methyltransferases DNMT3A and DNMT3B, and is maintained through cell division by the action of DNMT1 at hemi-methylated sites (Reik, 2007). Recent studies have explored the abundance of methylation at CH sites. Unlike plants, which have specific DNA methyltransferases that target CHG and CHH sites (Wendte and Schmitz, 2018), mCH in mammals relies on ‘off-target’ activity by DNMT3A and DNMT3B (Gabel et al., 2015; Guo et al., 2014; Lister et al., 2013; Ramsahoye et al., 2000; Stroud et al., 2017). There appears to be no mechanism to maintain asymmetrical CH methylation, however, which means that CH methylation is lost in replicating cells (He and Ecker, 2015). Post-mitotic neurons are unique among mammalian somatic cell types in that they accumulate high levels of CH methylation, most prevalently in CAC trinucleotides (Guo et al., 2014; Varley et al., 2013; Xie et al., 2012). This modification arises postnatally due to high levels of DNMT3A, which persist throughout adulthood (Feng et al., 2005; Stroud et al., 2017). The discovery that MeCP2 targets mCAC sites preferentially over all other forms of mCH (Gabel et al., 2015; Lagger et al., 2017; Liu et al., 2018; Sperlazza et al., 2017) raised the possibility that this DNA binding specificity contributes to neuronal function. Non-CG methylation almost doubles the number of available MeCP2 binding sites in neurons (Lagger et al., 2017; Lister et al., 2013; Mo et al., 2015) and the timing of increased CAC methylation coincides with the increase in neuronal MeCP2 protein levels during the first few weeks of life (Guo et al., 2014; Lister et al., 2013; Skene et al., 2010; Stroud et al., 2017).

It is difficult to disentangle the influence of mCG vs mCAC on regulation of gene expression by MeCP2, as these marks are interspersed with one another (Guo et al., 2014; Lavery et al., 2020). In the present study, we have assessed their relative importance using molecular genetic approaches to separate the effects of mCG and mCAC on MeCP2 function. Firstly, we directly visualised footprints of MeCP2 bound to mCG and mCAC in native brain chromatin. We also detected a subtle footprint over the rarer motif, mCAT, which had previously been reported to bind weakly to MeCP2 *in vitro* (Lagger et al., 2017). To determine the biological importance of the ability of MeCP2 to bind mCAC sites, we took advantage of the fact that the conserved methyl-CpG binding domain (MBD) of the related protein, MBD2, confers binding to mCG sites only. This allowed us to create a chimeric MeCP2-MBD2 (MM2) protein by domain swapping. Despite retention of mCG binding, knock-in mice expressing MM2 developed severe phenotypic features that largely mirrored those seen in mouse models of RTT. We conclude that binding to mCG alone is insufficient for MeCP2 to fulfil its role in the maintenance of neurological function. At a molecular level, inability to bind mCAC leads to global transcriptional changes in *MM2* mice, with a third of dysregulated genes also altered in *Mecp2*-null mice. Intriguingly, this shared subset is mostly upregulated, highly methylated and enriched for genes associated with neurological disease, highlighting potentially important contributors to the RTT-like phenotype.

## RESULTS

### MeCP2 footprints in native brain chromatin

MeCP2 binding to mCG and mCAC sites has been characterised *in vitro* using multiple techniques and the co-crystal structures of both these interactions have been solved (Ho et al., 2008; Lagger et al., 2017; Lei et al., 2019; Lewis et al., 1992; Mellén et al., 2012; Sperlazza et al., 2017). A third potential MeCP2 binding motif, mCAT, has been proposed, though this interaction appears to be weaker and is barely detectable in some assays (Lagger et al., 2017; Liu et al., 2018). We used BLI (Bio-Layer Interferometry) to quantify the interactions between the MBD of MeCP2 and DNA probes containing each of three methylated motifs (Figure 1A). MeCP2 binds to both mCG and mCAC, with dissociation constants (K_D_) of 22.25 ± 3.12 nM and 13.90 ± 0.64 nM, respectively. In contrast, binding to mCAT is ~3–5-fold weaker, with a K_D_ 63.17 ± 6.94 nM. Previously, full-length MeCP2 was shown to bind readily to all three sequences when co-overexpressed with methylated oligonucleotides in HEK293 cells (Lagger et al., 2017). To assess this further, we used beads coated with double-stranded oligonucleotides to pull-down endogenous MeCP2 from rat brain lysates. Like mCG and mCAC (Connelly et al., 2020), mCAT-containing DNA efficiently enriched MeCP2 protein in a DNA methylation-dependent manner (Figure 1B). Defining MeCP2 binding sites in native brain chromatin has proven more challenging due to the high abundance of both MeCP2 protein and its short target sequences. This is further complicated at mCH sites as the proportion of methylation at the vast majority of these loci is very low, in contrast to CGs that are highly methylated in the bulk genome (Figure 1C). ChIP-seq analysis produces a broad but relatively featureless signal across the genome, except at non-methylated CpG islands where the number of reads drops sharply (Chen et al., 2015; Kinde et al., 2016; Lagger et al., 2017; Skene et al., 2010). In spite of this limitation, peak-calling algorithms detect enrichment of mCG and mCAC (but not mCAT) coincident with the summits of MeCP2 binding (Gabel et al., 2015; Lagger et al., 2017). To assay genomic binding more robustly, we adopted the Assay for Transposase-Accessible Chromatin using sequencing (ATAC-seq) to reveal MeCP2-specific footprints. By dividing the ATAC-seq signal from wild-type samples by the equivalent signal from *Mecp2*-null (KO) samples, the method has revealed protected genomic regions attributable to MeCP2 at mCG in cultured human neurons (Cholewa-Waclaw et al., 2019). Here, we show that a MeCP2 binding footprint is also detectable over methylated CG dinucleotides in mouse hypothalamus tissue (Figure 1D). To detect a MeCP2 binding footprint at mCAC sites with this method, we focused on the subset of sites with >75% methylation (Figure 1E, S1A). The greater prominence of the mCAC footprint compared to the mCG footprint in hypothalamus cannot be interpreted as stronger binding to the trinucleotide motif, as footprint dimensions are affected to an unknown extent by several factors, including selection criteria for sites and sequence context (for example, CAC repeats, which are prevalent in the genome and can be methylated). Footprints were not initially detectable at mCAT, but by lowering stringency to include sites >50% methylated (Figure S1A), we detected a subtle footprint at mCAT, but not at other mC-containing motifs (Figure 1F, S1B-E). No footprints were detected over unmethylated cytosine in any sequence context (Figure 1D-F, S1B-E). We conclude that the main binding sites for MeCP2 are mCG and mCAC. Although interaction with mCAT moieties is detectable *in vitro* and *in vivo*, it is likely to be of less biological significance due to the reduced binding affinity and lower abundance of this motif.

**Figure 1.**
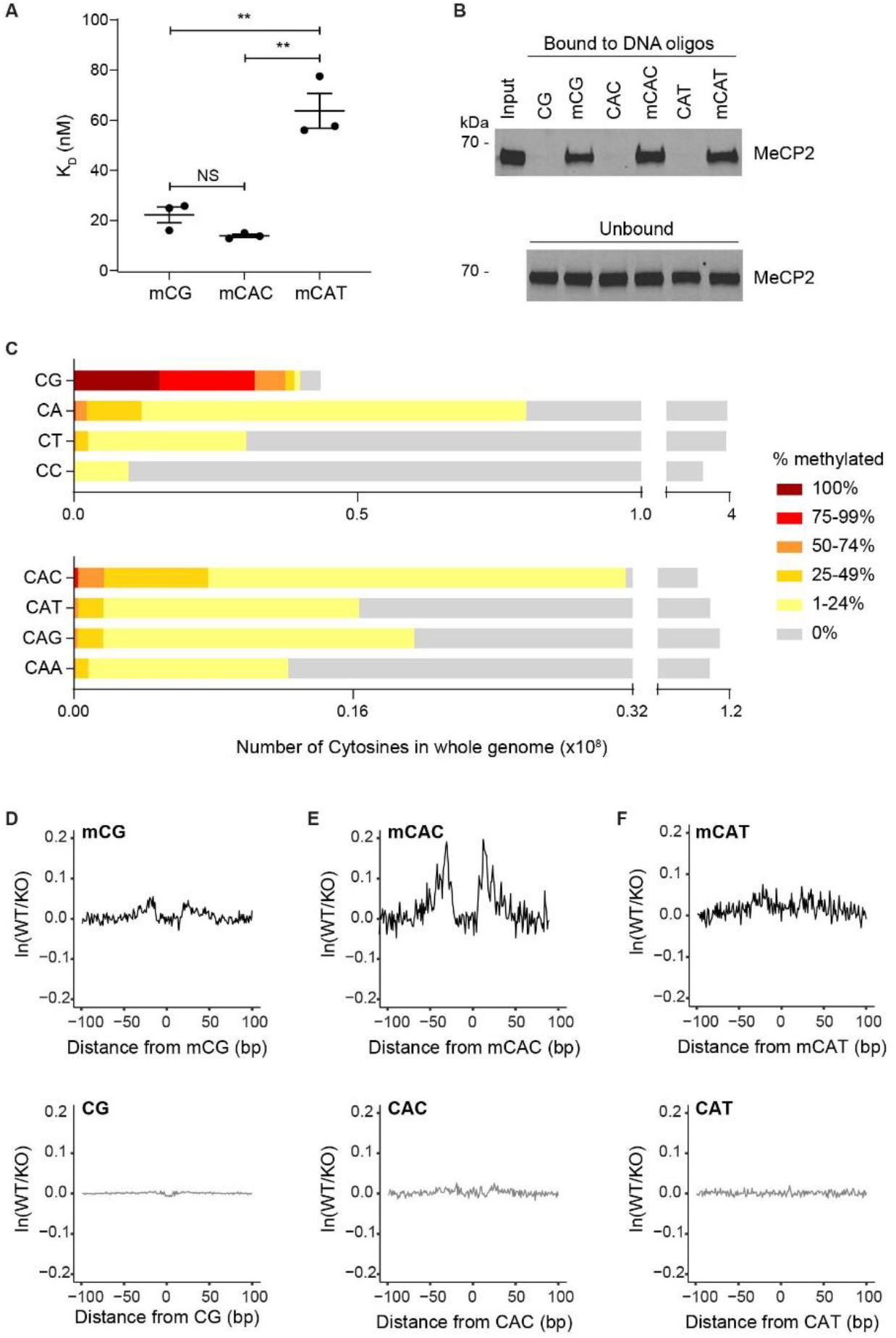
MeCP2 binds mCG and mCAC, and to a lesser extent, mCAT. A. Bio-Layer Interferometry (BLI) analysis of the interaction between the MBD of MeCP2 (residues 77-167) and methylated DNA probes containing a single mCG, mCAC or mCAT site. K_D_ values: mCG 22.25 ± 3.12 nM; mCAC 13.90 ± 0.64 nM; mCAT 63.71 ± 6.94 nM (mean ± S.E.M.). Binding affinity to mCG and mCAC is not significantly different (NS P = 0.06, t-test). Binding to mCAT is significantly weaker than mCG and mCAC (** P = 0.006 and ** P = 0.002, respectively, t-tests). B. Western blot analysis of MeCP2 following pull-down from rat brain nuclear extracts using immobilised DNA sequences containing unmethylated and methylated CG, CAC and CAT (upper) and unbound MeCP2 in the supernatant (lower). C. Whole genome bisulphite sequencing analysis in the mouse hypothalamus (Lagger et al., 2017) showing the distribution of methylation at each CN dinucleotide (upper) and each CAN trinucleotide (lower). Sites were binned by level of methylation. D-F. ATAC-seq footprinting analysis of MeCP2 over methylated (upper) and unmethylated (lower) CG (D), CAC (E) and CAT (F) sites in the mouse hypothalamus.

### A chimeric MeCP2 that selectively binds mCG

To determine the biological importance of the dual binding specificity of MeCP2, we sought to modify MeCP2 protein so that it could bind to only one of these motifs. A DNA binding domain that exclusively binds mCAC is unavailable, but domains that only bind mCG are present in other members of the MBD protein family. In particular, the evolutionarily most ancient member of the MBD family, MBD2 (Hendrich and Bird, 1998; Tweedie et al., 1999) can only bind to methylated cytosine in the canonical mCG context (Guo et al., 2014; Liu et al., 2018). We therefore substituted the MeCP2 MBD (94-164 aa) with the equivalent domain from MBD2 (amino acids 153 – 217) to create a chimeric “MM2” protein (Figures 2A-B). After confirming by EMSA analysis that the MBD of MBD2 readily binds to mCG only (Figure 2C), we performed similar analysis using N-terminal fragments of MM2 and WT MeCP2. As expected, MM2 adopted the binding specificity of MBD2, showing affinity for the mCG-containing but not the mCAC-containing DNA probe (Figure 2D).

**Figure 2.**
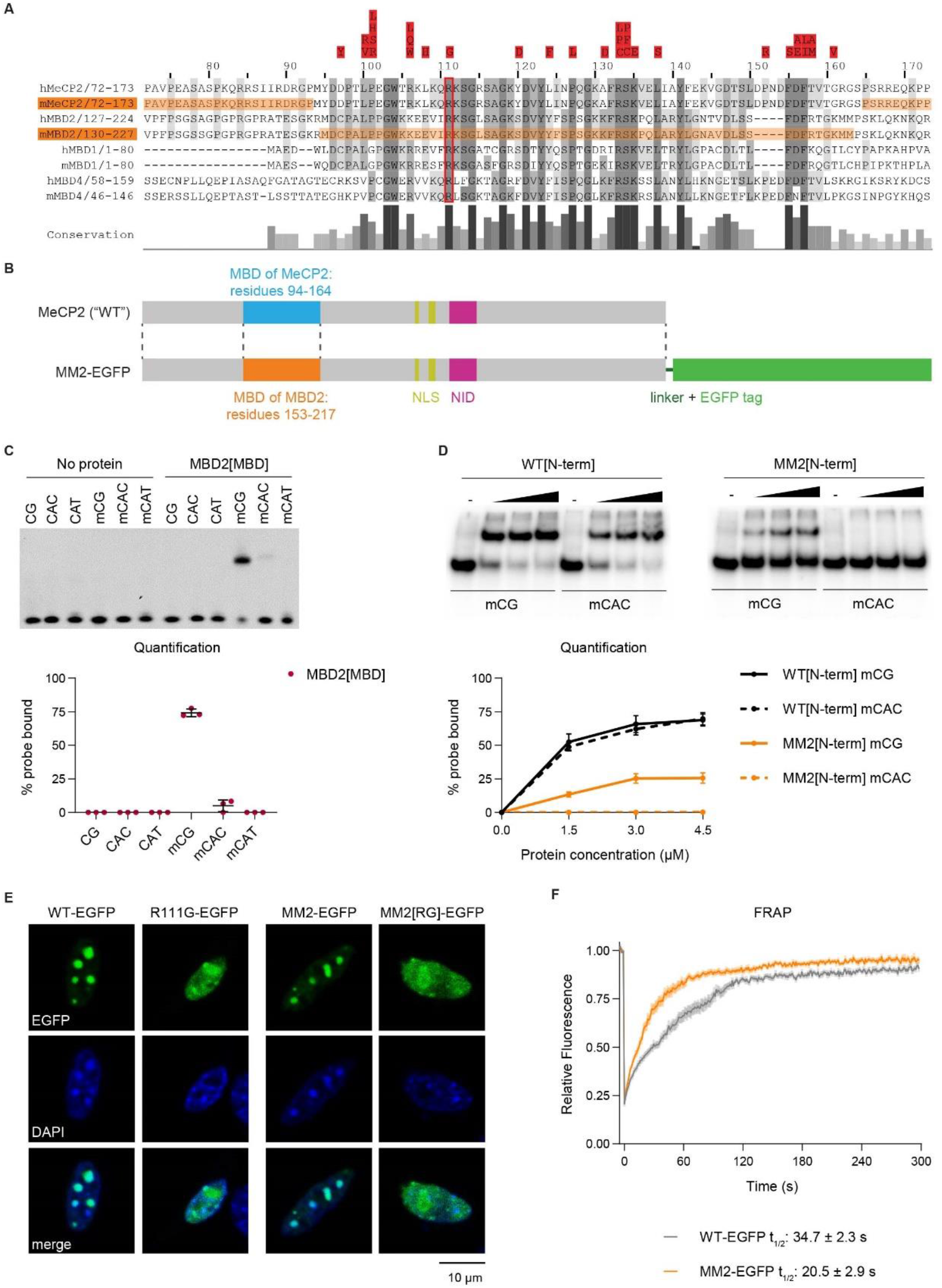
Chimeric protein MM2 has the DNA binding properties of MBD2 in the context of MeCP2. A. Alignment of human (h) and mouse (m) MeCP2, MBD2, MBD1 and MBD4, shaded by percentage identity, with conservation below. The RTT-causing missense mutations that lie in the MBD are shown in red (above). The conserved arginine at position 111 in MeCP2 is boxed in red. The sequences from mMeCP2 and mMBD2 used in MM2 are shaded in orange. B. Schematic showing the design of the chimeric protein MM2, where the MBD of MBD2 (residues 153-217) was inserted into the MeCP2 protein sequence to replace the endogenous MBD (residues 94-164). MM2 was tagged at the C-terminus with EGFP, connected with a short peptide linker. C. EMSA analysis of the MBD of human MBD2 binding to DNA probes containing a single unmethylated or methylated CG, CAC or CAT site. MBD2[MBD] protein concentration: 2 μM. Quantification is shown below. D. EMSA analysis of the binding affinities of N-terminal fragments (residues 1-205) of WT MeCP2 and MM2 to DNA probes containing a single mCG or mCAC site. The concentrations of each protein range from 0-4.5 μM. Quantification is shown below. E. Representative images showing localization of EGFP-tagged MeCP2 (WT) and MM2 proteins to mCG-rich heterochromatic foci when they are transiently overexpressed in mouse fibroblasts (NIH-3T3). Mutation of the MBD (R111G and the equivalent mutation in MM2) abolished binding, resulting in diffuse localisation. Scale bar, 10 μm. F. FRAP analysis at heterochromatic foci of WT-EGFP and MM2-EGFP. Total numbers of cells analysed: WT-EGFP, n = 27; MM2-EGFP, n = 28; over three independent transfection experiments. Graph shows fluorescence relative to prebleach, mean ± S.E.M. Half lives: WT-EGFP = 34.66 ± 2.32 s; MM2-EGFP = 20.46 ± 2.88 s (mean ± S.E.M.). Half-lives were compared using a Mann-Whitney test: **** P < 0.0001.

To test the ability of the chimeric protein to bind mCG sites *in vivo*, we transiently over-expressed the EGFP-tagged MM2 in cultured mouse fibroblasts, where mCG is highly concentrated in heterochromatic foci (Nan et al., 1996). MM2-EGFP co-localised efficiently with pericentromeric heterochromatin in a manner indistinguishable from WT-EGFP (Figure 2E). Mutating the essential arginine R111 (Figure 2A) to glycine disrupted the localisation of MeCP2 (Figure 2E) (Kudo et al., 2003) and the equivalent mutation in MM2-EGFP had the same effect (Figure 2E), confirming that mCG binding is mediated by this MBD. To assess the binding dynamics between MM2-EGFP and heterochromatin, we performed transient transfection followed by Fluorescence Recovery After Photobleaching (FRAP). WT-EGFP binding recovered at heterochromatic foci with a half-life of 34.7 ± 2.3 s (Figure 2F), consistent with previous studies (Ghosh et al., 2010; Klose et al., 2005; Ludwig et al., 2017). The half-life of MM2-EGFP was 20.5 ± 2.9 s, indicating that molecules of the chimeric protein exchange slightly faster (P < 0.0001, Mann-Whitney test). This may reflect a somewhat reduced binding affinity for mCG, as also suggested by the lower percentage of probe bound in the EMSA assay (Figure 2D). Notably, this difference is much smaller than the destructive effect of RTT-causing mutations that lie in the MBD domain (Ballestar et al., 2000; Schmiedeberg et al., 2009).

The key function of MeCP2 is to form a bridge between methylated DNA and the NCOR1/2 corepressor complexes, dependent on the MBD and NID (NCOR1/2 Interaction Domain), respectively (Guy et al., 2018; Kruusvee et al., 2017; Lyst et al., 2013; Tillotson et al., 2017). MM2 retains the NID (residues 298-309) and accordingly was able to pull-down NCOR1/2 complex subunits HDAC3 and NCOR1 when transiently over-expressed in HeLa cells (Figure S2A). Likewise, MM2 retains the ability to recruit TBL1X (its direct binding partner in the NCOR1/2 complexes) to mCG-rich heterochromatic foci when co-expressed in cultured fibroblasts (Figure S2B). The NID-abolishing RTT mutation, R306C, served as a negative control in both these assays (Lyst et al., 2013). Overall, the behaviour of MM2 in all *in vitro* and cell culture-based assays supports the notion that this chimeric protein retains the essential properties of MeCP2, with the major exception that its DNA binding is restricted to mCG sites.

### MM2 selectively binds mCG *in vivo* in knock-in mice

To determine the function of MM2 *in vivo*, we produced knock-in mice by replacing the endogenous *Mecp2* allele with DNA sequence encoding *MM2* fused to a C-terminal EGFP tag (Figure S3). Quantification of MM2-EGFP protein expression in mouse brain using western blotting and flow cytometry demonstrated approximately equivalent levels when compared to mice with WT-EGFP (Figure 3A-B), which is itself expressed at the same level as the native untagged protein (Tillotson et al., 2017). The localisation of MM2-EGFP at mCG-rich pericentromeric heterochromatin in the brain mimics WT-EGFP, indicative of mCG binding *in vivo* (Figure 3C). We used ChIP-qPCR to quantify mCG binding with primers annealing to genes in the relatively mCA-poor cytochrome p450 locus (Brown et al., 2016). Binding of MM2-EGFP was comparable to WT-EGFP at all three genes (Figure 3D), indicating that MM2-EGFP binds mCG *in vivo* in a similar manner to endogenous MeCP2. To assess the binding specificity of MM2-EGFP *in vivo*, we performed ATAC-seq on hypothalamus tissue, again dividing by the profiles derived from *Mecp2*-null (KO) tissue to reveal MM2-EGFP-dependent chromatin inaccessibility. A clear footprint was observed at mCG sites, but binding was undetectable at mCAC and mCAT motifs (Figure 3E-G). ATAC-seq analysis also showed that, like MeCP2, MM2-EGFP was unable to bind mC in other sequence contexts or to unmethylated cytosine (Figures 3E-G, S4). This result provides striking confirmation that the MBD domain swap has uncoupled the DNA binding specificities of MeCP2, abolishing binding to non-CG methylation.

**Figure 3:**
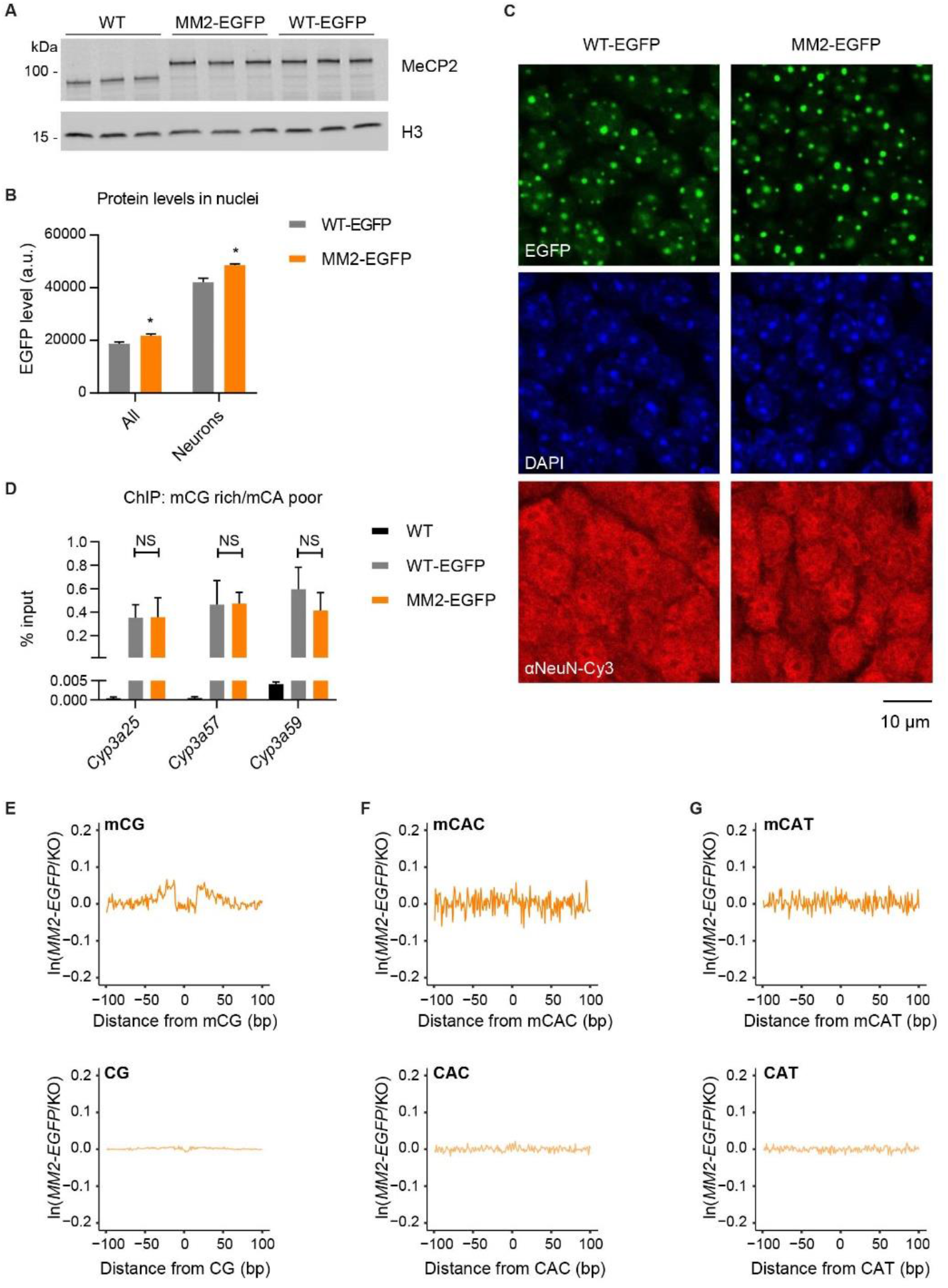
MM2 binds mCG but not mCAC or mCAT *in vivo*. A. Western blot analysis of whole-brain extract showing protein sizes and abundance of MM2-EGFP in whole brain compared to WT and WT-EGFP control, detected using an N-terminal MeCP2 antibody. Histone H3 was used as a loading control. B. Flow cytometry analysis of protein levels in nuclei from whole brain (All) and the high-NeuN subpopulation (Neurons) in WT-EGFP (n = 3) and MM2-EGFP (n = 3), detected using EGFP fluorescence. Graph shows mean ± S.E.M. and genotypes were compared by t-tests: All * P = 0.0.30; Neurons * P = 0.018; au, arbitrary units. C. WT-EGFP and MM2-EGFP are both localised at mCG-rich pericentromeric heterochromatin in the brains of knock-in mice. Representative images of the dentate gyrus of the hippocampus are shown. WT-EGFP and MM2-EGFP were visualised by EGFP fluorescence, heterochromatic foci appear as DAPI bright spots and neurons were stained with an αNeuN-Cy3 conjugate antibody. Scale bar = 10 μm. D. ChIP-qPCR analysis of WT-EGFP and MM2-EGFP at chromatin sites with high mCG and low mCAC (Brown et al., 2016). Chromatin was extracted from whole brain and pulled down with GFP-TRAP beads. Chromatin from WT (untagged) brain samples was used at a negative control. There were similar levels of WT-EGFP and MM2-EGFP at all three loci, compared using t-tests (all NS, P > 0.05). E-G. ATAC-seq footprinting of MM2-EGFP at methylated (upper) and unmethylated (lower) CG (E), CAC (F) and CAT (G) sites. ATAC-seq profiles from the hypothalamus of *MM2-EGFP* mice were normalised to *Mecp2*-null (KO) profiles at each site. Equivalent WT ATAC-seq footprinting is shown in Figures 1D-F.

### MM2 knock-in mice display a Rett syndrome-like phenotype

To determine the phenotypic consequences of the domain-swap, we assessed weekly a cohort of hemizygous male *MM2-EGFP* mice and wild-type littermate controls (n = 10 per genotype) for overt symptoms associated with mouse models of RTT: hypoactivity, abnormal gait, hind-limb clasping, tremor, irregular breathing and deterioration of general condition (Guy et al., 2007). Like mice with RTT-causing mutations, *MM2-EGFP* mice were born healthy and onset of symptoms occurred shortly after weaning (Figure 4A). The phenotypes became progressively more severe until animals reached their humane endpoint, with a median survival of 29.5 weeks (Figure 4B). *MM2-EGFP* mice weighed less than their wild-type littermates throughout their lifetime (Figure 4C), consistent with RTT models on a C57BL/6J background (Brown et al., 2016; Guy et al., 2001). Additionally, acute loss of bodyweight, which accompanies severe disease in RTT models, was correlated with morbidity in the *MM2-EGFP* mice. Mouse models of RTT vary in severity depending on MeCP2 mutation, mirroring trends observed in patients (Brown et al., 2016; Cuddapah et al., 2014). *MM2-EGFP* mice lie in the middle of the RTT severity spectrum, closely following the progression seen in *R306C-EGFP* mutant mice (Figures 4D-F). More detailed analysis of the six overt symptoms that were scored weekly revealed that *MM2-EGFP* mice recapitulated the RTT phenotypic signature, with the exception of tremors, which were not apparent (Figure 4G). This was most evident when phenotypic severity scores of *MM2-EGFP, R306C-EGFP* and *R133C-EGFP* mice were aligned to focus on the five weeks prior to reaching their humane endpoint (Figure 4H).

**Figure 4:**
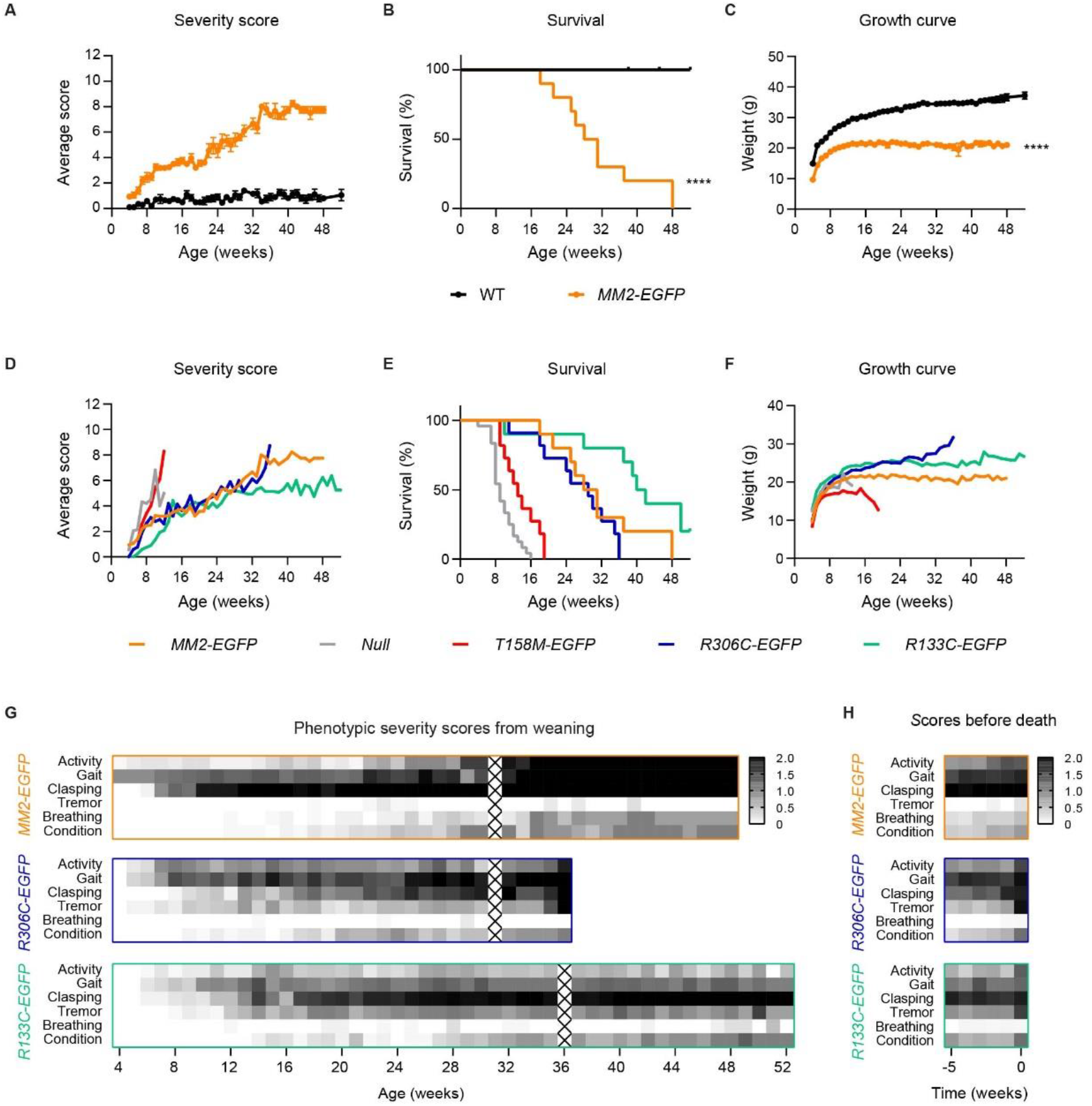
*MM2* knock-in mice display overt RTT-like neurological defects. A. Phenotypic severity scores of hemizygous male *MM2-EGFP* mice (n = 10) compared with their WT littermates (n = 10) over one year. Graphs show mean scores ± S.E.M. B. Kaplan–Meier plot of survival of the scoring cohort. Two wild-type controls were culled due to injuries, aged 38 and 45 weeks (censored). Genotypes were compared using a Mantel–Cox test: **** P < 0.0001. C. Growth curve of the scoring cohort. Graph shows mean values ± S.E.M. Genotypes were compared using Mixed-effects analysis: **** P < 0.0001. D-F. Phenotypic severity score (D), survival (E) and growth curves (F) of *MM2-EGFP* mice over 1 year compared to *Mecp2*-null mice (n = 12/24/20) and models carrying patient mutations (all with a C-terminal EGFP tag): *T158M* (n = 11/11/15), *R306C* (n = 11, same cohort for all) and *R133C* (n = 10, same cohort from all) (Brown et al., 2016). G-H. Heatmaps of the phenotypic scores of *MM2-EGFP*, *R306C-EGFP* and *R133C-EGFP* mice shown in (D), divided into the six categories. The plots are shaded according to the mean score for each category. Scores were analysed according to age (G) and for the 5 weeks prior to each mouse reaching its humane end-point (H). Weeks where mice were not assessed are marked with an X.

A second cohort of *MM2-EGFP* male mice and wild-type littermates (n = 10 per genotype) underwent a series of behavioural tests at 10-11 weeks of age. Like RTT models (Brown et al., 2016), *MM2-EGFP* mice displayed decreased anxiety in the elevated plus maze, spending significantly more time in the open arms than wild-type littermate controls (Figure 5A). *MM2* mice were slightly hyperactive in the open field test (Figure 5B), which differs from the reduced activity characteristic of RTT models. However, both *MM2-EGFP* and RTT models displayed progressive loss of spontaneous activity as phenotypes worsened (Figure 4G-H). *MM2-EGFP* mice exhibited a trend towards reduced motor coordination in the hanging wire test (Figure 5C). Motor function was better assessed by the accelerating rotarod, where wild-type mice showed increasing ability to stay on the rotarod over the three days, but the *MM2-EGFP* mice had significantly impaired performance compared to the controls on days two and three (Figure 5D). Motor deficits are observed in all mouse models of RTT (Brown et al., 2016; Goffin et al., 2012; Tillotson and Bird, 2019). In summary, *MM2-EGFP* mice share most of the phenotypic features found in mice that are deficient in functional MeCP2, including delayed symptom onset, hind limb clasping, abnormal gait, motor defects, and decreased anxiety.

**Figure 5:**
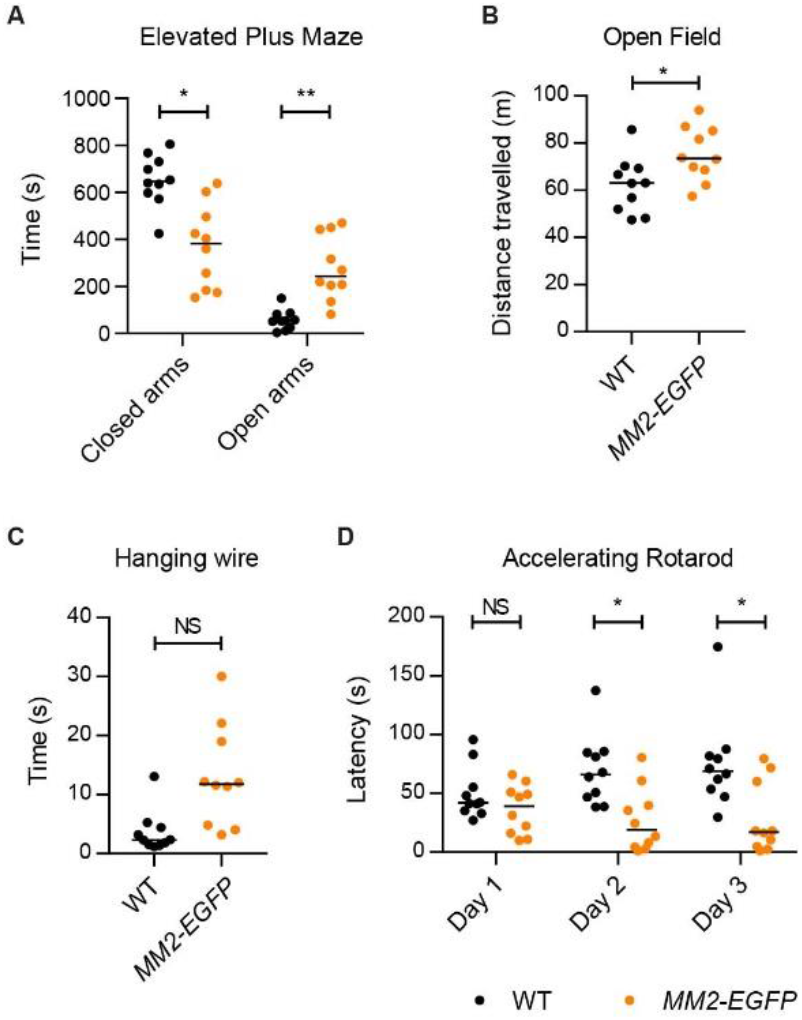
MM2 knock-in mice display behavioural defects associated with the RTT-like phenotype. A-D. Behavioural analysis of a separate cohort performed at 10-11 weeks of age: *MM2-EGFP* (n = 10), compared with their WT littermates (n = 10). Graphs show individual values and medians, and statistical significance as follows: not significant (NS), P > 0.05, * P < 0.05, ** P < 0.01. A. Time spent in the closed and open arms of the elevated plus maze during a 15 min trial. Genotypes were compared using Kolmogorov–Smirnov tests: closed arms, * P = 0.015; and open arms, ** P = 0.003. B. Distance travelled in open field test was measured during a 20 min trial. Genotypes were compared using a t-test: * P = 0.022. C. Mean time taken for animals to bring a hind paw to the wire in the hanging wire test in three trials (each up to 30 s). Genotypes were compared using a KS test: P = 0.055. D. Mean latency to fall from the accelerating rotarod in four trials was calculated for each of the 3 days of the experiment. Genotypes were compared using Kolmogorov–Smirnov tests: day 1, P = 0.401; day 2, * P = 0.015; and day 3, * P = 0.015. WT animals displayed significant improvement over the three days (* P = 0.026), *MM2-EGFP* animals did not change significantly (P = 0.314), analysed with Friedman tests.

### MM2 represses transcription at mCG but not mCAC sites

Global transcriptional changes caused by deficiency of functional MeCP2 are thought to underlie RTT (Chahrour et al., 2008; Chen et al., 2015; Gabel et al., 2015; Kinde et al., 2016; Lagger et al., 2017; Renthal et al., 2018). These studies have demonstrated that transcriptional upregulation correlates with the level of methylation, consistent with the role of MeCP2 in conferring repression at methylated sites. To explore the role of the two different motifs, we performed RNA-seq on hypothalamus tissue harvested from *Mecp2*-null (KO) mice and wild-type controls at 6 weeks of age, when KO mice are symptomatic but no animals had reached their humane endpoint. To facilitate comparison between mutants, tissue from *MM2-EGFP* mice and wild-type littermate controls was harvested at both 6 weeks (age-matched with KO) and 12 weeks (symptom-matched with KO). As expected, transcriptional changes in KO/WT were positively correlated with the number of MeCP2 binding sites (mCG + mCAC) per gene (Figure 6A). Similar trends were also seen in the *MM2-EGFP*/WT datasets (Figure 6B, S5A). Surprisingly, when we compared transcriptional changes to the number of mCG or mCAC binding sites alone, positive slopes were observed for both motifs in the *MM2-EGFP*/WT datasets as well as KO/WT (Figure 6A-B, S5A). This gives the impression that mCG-dependent repression is reduced in *MM2-EGFP* mice, but can be explained by the tight correlation between the number of mCG and mCAC binding sites in each gene (Figure 6C, r = 0.90).

**Figure 6.**
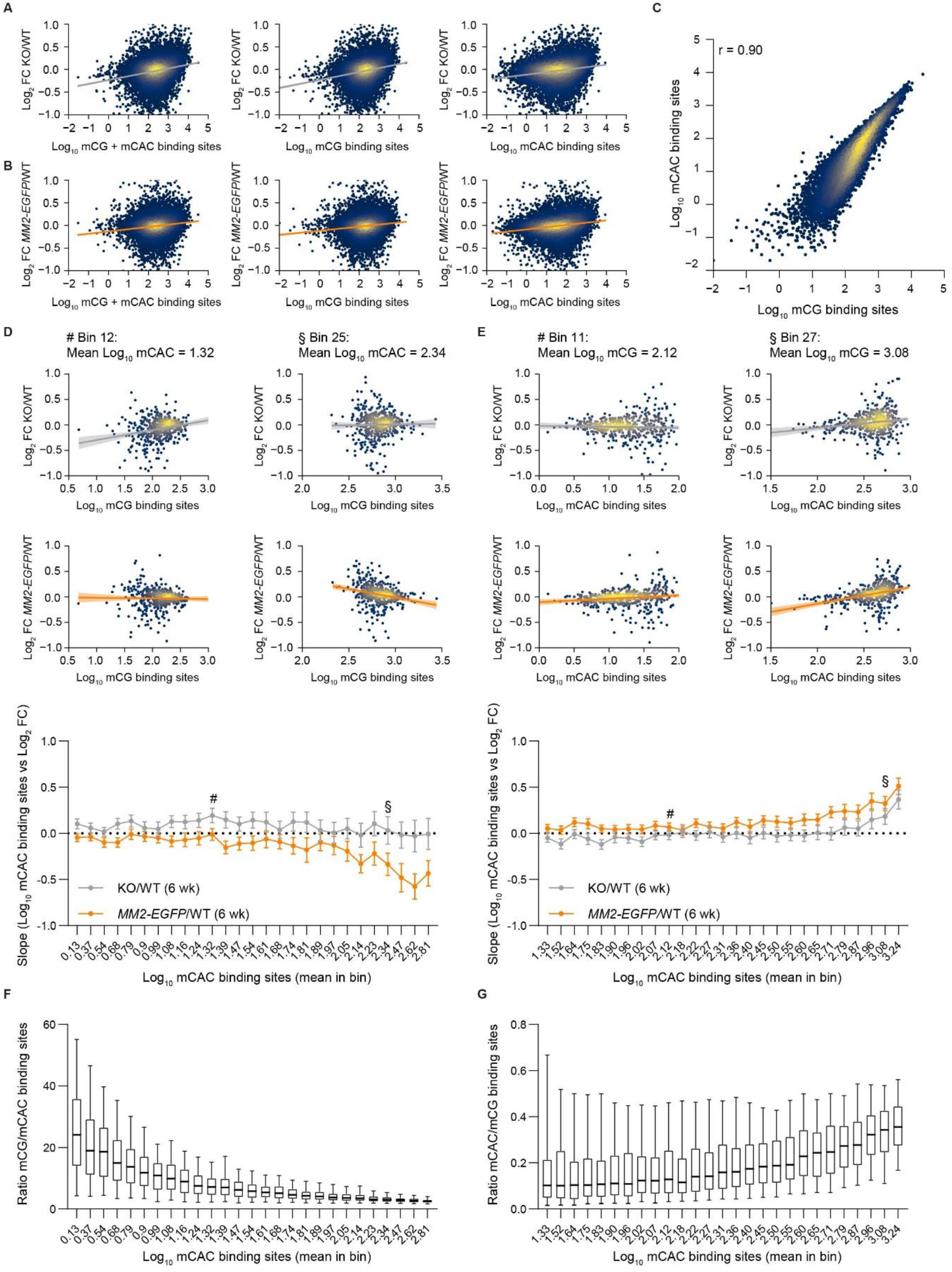
MM2 represses transcription at mCG but not mCAC sites. A-B. Correlations between the total number of MeCP2 binding sites per gene: mCG + mCAC (left), mCG (centre) and mCAC (right) and transcriptional changes in KO/WT (A) and *MM2-EGFP*/WT (B) hypothalamus tissue at 6 weeks of age. C. Correlation between mCG and mCAC binding sites in all genes in the hypothalamus at 6 weeks (measured as log_10_ sites per gene). Regression (r) = 0.90. D. Genes were binned by the number of mCAC binding sites to determine the effect of mCG on transcription in KO/WT and *MM2-EGFP-EGFP/WT* (6 weeks). Two example bins are shown (# Bin 12, mean log_10_ mCAC = 1.32; and § Bin 25, mean log_10_ mCAC = 2.34). The slopes of all bins are shown below. E. Genes were binned by the number of mCG binding sites to determine the effect of mCAC on transcription. Two example bins are shown (# Bin 11, mean log_10_ mCG = 2.12; and § Bin 27, mean log_10_ mCAC = 3.08). The slopes of all bins are shown below. Error on all slopes: 95% CI. F. The ratio of mCG/mCAC binding sites per gene in each mCAC bin. G. The ratio of mCAC/mCG binding sites per gene in each mCG bin.

To uncouple the effects of the two motifs, we binned genes by one motif to determine the effect of the other. We predicted that transcriptional changes in KO relative to WT would positively correlate with both mCG and mCAC levels. Surprisingly, positive correlations between mCG levels and transcriptional changes were only present in bins with lower methylation, with highly methylated genes showing no correlation (Figure 6D). In contrast, positive trends between mCAC levels and KO/WT transcription changes increased as genes became more methylated (Figure 6E). Interestingly, the proportion of methylated sites per gene that are within mCAC motifs increases with total methylation (Figure 6F-G). Our results reveal that mCAC sites have a greater impact on the regulation of more methylated genes (where mCAC/mCG > 0.3). We predicted that transcriptional changes in *MM2-EGFP* mice compared to wild-type controls would show a positive correlation with mCAC but not mCG. Indeed, there was no correlation between mCG levels and *MM2-EGFP*/WT transcriptional changes in lowly methylated genes (Figure 6D, S5B), consistent with MM2-EGFP retaining the ability to target mCG sites. In bins containing genes with higher methylation, we observed negative correlations between mCG level and *MM2-EGFP*/WT transcriptional changes, suggesting greater mCG-dependent repression in *MM2-EGFP* hypothalamus compared to wild-type controls. A possible explanation is higher MM2-EGFP occupancy at mCG sites, due to its inability to target mCAC sites resulting in more free molecules. Alternatively, as mCG levels increase within each bin, the proportion of retained MM2-EGFP binding sites (in other words, the mCG/mCAC ratio) per gene increases (Figure S5C). Analysis of mCAC trends showed a similar pattern for *MM2-EGFP*/WT as for KO/WT (Figure 6E, S5D), consistent with loss of mCAC-dependent repression in both mutants. The positive slopes were slightly steeper in the *MM2-EGFP*/WT datasets, explained by the link between mCAC levels and the proportion of lost MM2-EGFP binding sites (i.e. mCAC/mCG ratio) (Figure S5E). In summary, MM2-EGFP protein has specifically lost the ability to repress transcription at mCAC sites, disproportionally affecting genes with higher overall levels of methylation.

### Shared dysregulated genes are implicated in disease

RTT may be caused by the aggregate effect of small transcriptional changes at hundreds of genes. Alternatively, a few dysregulated genes may be primarily responsible for the phenotypes, the remainder being relatively neutral. Given the remarkable similarity between the phenotypes of *MM2-EGFP* and RTT mice, these genes would be affected in both mutants. Significantly changed genes overlapped by about a third in both the age-matched and symptom-matched datasets (Figure 7A, purple shading). Importantly, for almost all of the 316 shared genes, the direction of transcriptional change was the same in KO and *MM2-EGFP* mice (Figure 7B, S6A). To explore the possibility that the shared genes could lead us to candidates whose altered expression contributes strongly to the RTT phenotype, we performed Disease Ontology analysis on each of the three groups: shared (purple), KO only (grey) and MM2 only (orange). We found that the shared genes were enriched in genes associated with four categories of neurological disease: “pervasive developmental disorder”, “autism spectrum disorder”, “autistic disorder” and “developmental disorder of mental health” (Table S1). It is notable that the same analysis on genes dysregulated only in KO mice or only in *MM2-EGFP* mice gave no significant disease hits. There was a strong overlap between the four neurological disease categories, with all 20 genes associated with “developmental disorder of mental health” present in one or more of the other categories. The candidate genes were similarly dysregulated in both mutants in this study and an independent hypothalamus dataset (Figure 7C) (Chen et al., 2015). Many of these genes were also dysregulated in KO cortical tissue (Figure 7C) (Boxer et al., 2019), suggesting consistency between different regions of the RTT brain.

**Figure 7.**
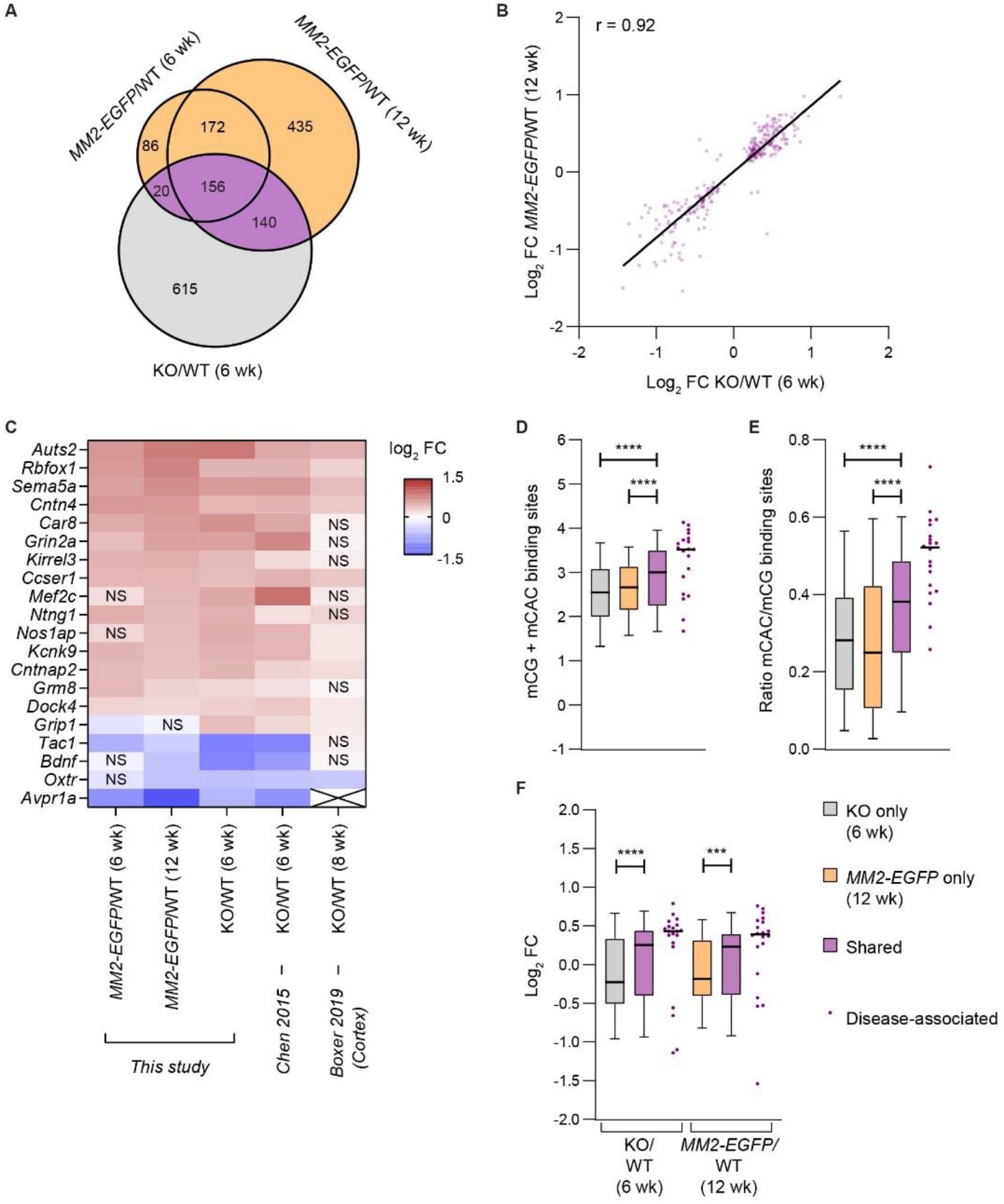
Shared dysregulated genes are linked to neurological disease. A. Venn diagram showing the overlap between significantly dysregulated genes in KO (6 weeks), *MM2-EGFP* (6 weeks) and *MM2-EGFP* (12 weeks), each compared to aged-matched wild-type controls. Significance cut-off: padj < 0.05. B. Shared genes are dysregulated in the same direction in KO/WT (6 weeks) and *MM2-EGFP*/WT (12 weeks). C. Heatmap showing transcriptional changes in the 20 neurological disease-associated genes. The three datasets in this study were compared to an independent KO/WT (6 weeks) hypothalamus dataset (Chen et al., 2015) and a KO/WT (8 weeks) cortex dataset (Boxer et al., 2019). NS = not significant (padj > 0.05). *Avpr1a* expression is not present in the cortex dataset, denoted by the X. D. Shared genes have more mCG + mCAC binding sites, compared to genes dysregulated in only KO or only *MM2-EGFP* (12 weeks). E. Shared genes have higher mCAC/mCG ratios, compared to genes dysregulated in only KO or only *MM2-EGFP* (12 weeks). F. Shared genes are more upregulated in both mutants, compared to genes dysregulated in only KO or only *MM2-EGFP* (12 weeks). D-F. Disease-associated genes have exceptionally high mCG + mCAC levels, mCAC/mCG ratios and are highly upregulated. Median values are indicated by a black bar. Pairs of gene sets were compared using Mann-Whitney tests: **** P < 0.0001, *** P < 0.001.

Differential gene expression could contribute to the disease state regardless of the direction of change, but 15 out of 20 candidate genes were upregulated in both mutants (Figure 7C), suggesting they are direct MeCP2 targets. Comparisons between all shared dysregulated genes and those affected in KO or *MM2-EGFP* only showed that shared genes had more methylated binding sites (mCG+mCAC) (Figure 7D, S6B), higher mCAC/mCG ratios (Figure 7E, S6C) and tended to be more highly upregulated than other significantly changed genes (Figure 7F, S6D). Interestingly, the 20 candidate genes have exceptionally high levels of methylation, proportion of mCAC sites and transcriptional upregulation (Figure 7D-F, S6B-D). These genes represent strong candidates for future investigation as their dysregulation may contribute disproportionately to the neurological features of RTT.

## DISCUSSION

This study investigated the biological significance of the dual binding specificity of MeCP2. MBD functionality is essential for the role of MeCP2 protein but, as RTT-causing missense mutations in this domain impact both mCG and mCAC binding (Brown et al., 2016), the importance of its ability to target mCAC sites was previously unknown. Our results show that mice dependent on a derivative of MeCP2 that can bind mCG but not mCAC develop severe RTT-like symptoms. We conclude that the ability of MeCP2 to target mCAC is essential to avoid the neurological problems associated with RTT and cannot be compensated by binding to mCG alone.

### The deposition and interpretation of non-CG methylation is essential for neurological function

Previous studies have investigated whether loss of the mCAC writer, *de novo* DNA methyltransferase 3A (DNMT3A), has similar consequences to loss of the reader, MeCP2 (Gabel et al., 2015; Lavery et al., 2020; Stroud et al., 2017). Although DNMT3A preferentially methylates CG over CH (Ramsahoye et al., 2000), its expression in postnatal neurons, particularly during the first three weeks of life (Feng et al., 2005), mainly leads to mCH (predominantly mCAC) deposition, as the majority of CG dinucleotides have already been methylated in early development (Lavery et al., 2020; Stroud et al., 2017). The difference in timing means that writing of these marks can be uncoupled by Cre-mediated deletion *of Dnmt3a* after the early embryonic developmental deposition of mCG but prior to the postnatal deposition of mCH. Accordingly, deletion of *Dnmt3a* in the central nervous system (CNS) between E7.5 and E15.5 using *Nestin-Cre* abolishes mCH in the brain with a much lower impact on mCG (Dubois et al., 2006; Gabel et al., 2015). These mice display neurological defects including hypoactivity, hind limb clasping, gait and motor defects, resulting in premature death at 18 weeks of age (Nguyen et al., 2007; Stroud et al., 2017). The strong overlap with the RTT phenotype is consistent with interpretation of this mark by MeCP2. At the molecular level, loss of either DNMT3A or MeCP2 results in overlapping transcriptional changes (Gabel et al., 2015; Lavery et al., 2020). Our *MM2* mice build on this evidence by showing that specifically abolishing the ability of MeCP2 to read mCH is sufficient to cause many of the neurological defects and transcriptional changes associated with the RTT phenotype.

It has been speculated that the postnatal deposition of mCH could explain the delayed onset of the RTT phenotype (Chen et al., 2015; Lavery et al., 2020), implying that MeCP2 binding to canonical mCG sites may be of lesser importance. Indeed, mCG binding is largely dispensable in early development (before symptom onset at weaning) and at all developmental stages in peripheral tissues where only mCG sites are available (Guy et al., 2001; Ross et al., 2016). In neurons, however, MeCP2 levels are much higher than any other cell type, plateauing at ~5 weeks of age at a level that allows MeCP2 to coat neuronal chromatin (Skene et al., 2010). In the absence of any evidence that MeCP2 interprets mCG and mCAC sites differently, the relative importance of these two motifs likely depends on their abundance and location. Considering first abundance, in mature neurons, the evidence suggests that mCG and mCH binding sites occur with approximately equal frequencies (Mo et al., 2015). It is therefore possible that the accumulation of mCAC simply increases the number of available MeCP2 binding sites. If so, replacing MeCP2 with MM2, which retains the ability to confer transcriptional repression at more than half of MeCP2 binding sites, would be equivalent to halving MeCP2 protein levels. Our findings rule out this possibility, as mice expressing MeCP2 at ~50% normal levels display only very mild behavioural phenotypes (Samaco et al., 2008), compared with the severe RTT-like symptoms seen in *MM2* mice.

It is therefore likely that differences in the patterning of mCG and mCAC sites underlie their distinct functional significance. CG dinucleotides are globally depleted in the genome, but highly methylated from an early stage of development. This means that the number of mCGs per gene approaches the number of CG dinucleotides and is relatively uniform between cell types. In contrast, the number of CH motifs is more than an order of magnitude higher than CG, but they become sparsely methylated in neurons (predominantly at CAC) after mCG patterns have already been laid down. Importantly, the application of mCAC methylation is not uniform, but targets genes in inverse proportion to their expression level during the early postnatal period (Stroud et al., 2017). We show here that, on average, highly methylated genes acquire a greater proportion of mCAC motifs, so are disproportionally affected by mCAC-dependant repression by MeCP2. Loss of mCAC-dependent repression results in their upregulation in both *Mecp2*-null and *MM2-EGFP* mice, suggesting that their dysregulation contributes substantially to neurological defects. Patterns of mCAC methylation have also been reported to be more cell type-specific, as a result of differences in expression during its deposition (Mo et al., 2015; Stroud et al., 2017). Future work will address the roles of neuronal subtype-specific patterns of mCAC-dependent transcriptional modulation of highly methylated genes for maintaining brain function.

### Candidate genes may be responsible for most RTT symptoms

Since the discovery that MeCP2 controls transcription, the RTT field has searched for key target genes. While some genes (e.g. *Bdnf*) have been shown to have larger effects (Chang et al., 2006), the preferred view is that the neurological condition is an aggregate consequence of subtle changes in hundreds of genes. Additionally, many of these genes are down-regulated, a likely consequence of smaller cell size (Lagger et al., 2017; Li et al., 2013; Yazdani et al., 2012). In this study, we focused on genes dysregulated in both mutants, given their overlapping phenotypes. This list comprises a third of genes altered in *Mecp2*-null mice (n = 136), which tend to be highly methylated and therefore direct MeCP2 targets. Of the 20 candidates identified by Disease Ontology analysis, 15 were markedly upregulated in both mutants. Neurological disease is often linked with haploinsufficiency of these genes, raising the possibility that dosage may be critical, with their overexpression also being deleterious, as is the case for *MECP2* (Van Esch et al., 2005). Genes implicated in neurological disease typically encode proteins falling into two categories: those with specific roles in neuronal function and chromatin-associated factors. The candidates identified in this study span both classes. For example, CNTN4 is a neuronal membrane glycoprotein (Oguro-Ando et al., 2017) and AUTS2 activates transcription in association with non-canonical PRC1 (Gao et al., 2014). Time will tell whether dysregulation of these candidates is involved in RTT pathology and whether any could be targeted therapeutically.

### Concluding remarks

It is interesting to speculate on the origin of the dual binding specificity of MeCP2. There is evidence that MeCP2 originally evolved as an mCG-specific DNA binding protein (Hendrich and Tweedie, 2003), as other members of the MBD protein family (MBD1, MBD2 and MBD4) cannot bind to mCH (Guo et al., 2014; Liu et al., 2018). We speculate that during the course of evolution, MeCP2 has added to its repertoire the ability to bind mCAC and, with lower affinity, mCAT. This is permitted by the relative flexibility of the sidechain of arginine 133, which is one of a pair of key arginines involved in DNA recognition (Ho et al., 2008; Lagger et al., 2017; Lei et al., 2019). In contrast, the equivalent arginine side-chain in MBD2 is constrained by interactions with other parts of the protein (Liu et al., 2018; Scarsdale et al., 2011). According to this view, the coevolution of DNMT3A and MeCP2 has turned two somewhat peripheral – perhaps originally biologically unimportant – properties of each protein (mCAC methylation and mCAC binding, respectively) into essential contributors to brain stability and function (see Bird, 2020).

## ACKNOWLEDGEMENTS

This work was supported by a Rett Syndrome Research Trust consortium grant, a Wellcome Centre grant (091580/Z/ 10/Z), a Wellcome Investigator Award (107930/Z/15/ Z) and a European Research Council grant (EC 694295 Gen-Epix) to A.B., a Ludwig Institute for Cancer Research grant, a Biological Sciences Research Council grant (BB/M001873/1) and a Conrad N. Hilton Foundation grant to S.K. R.T. was funded by a Biotechnology and Biological Sciences Research Council Doctoral Training Partnership studentship. We thank the following people for assistance: A. Cook (advice on designing the truncated proteins), S. Picaud (providing purified MBD2[MBD] protein), A. McClure (animal husbandry), M. Waterfall (flow cytometry) and A. Kerr (statistics). We also thank members of the Bird, Kriaucionis, M. E. Greenberg, G. Mandel and M. J. Justice laboratories for helpful discussions. A.B. and M. J. L. are members of the Simons Initiative for the Developing Brain at the University of Edinburgh. R.T. is currently a Sir Henry Wellcome Postdoctoral Fellow (210913/Z/18/Z).

All authors declare no competing interests.

## AUTHOR CONTRIBUTIONS

Conceptualization, A.B., R.T., M.J.L. and S.K.; Methodology, R.T., J.C.W., S.A.K. and S.K.; Software, K.C., S.W. and D.K.; Formal analysis, K.C., S.W., R.T., J.C., and S.A.K.; Investigation, R.T., J.C.W., S.A.K., J.C., M.V.K., J.S., D.D.S. and K.B.; Writing – Original Draft, A.B. and R.T.; Writing – Review & Editing, all authors; Visualization, R.T., K.C., S.W., S.A.K. and J.C.; Data curation, K.C., S.W. and D.K.; Supervision, A.B., and S.K.; Project administration, J.S. and D.D.S; Funding Acquisition, A.B., and S.K.

## SUPPLEMENTAL FIGURES AND LEGENDS

**Figure S1.**
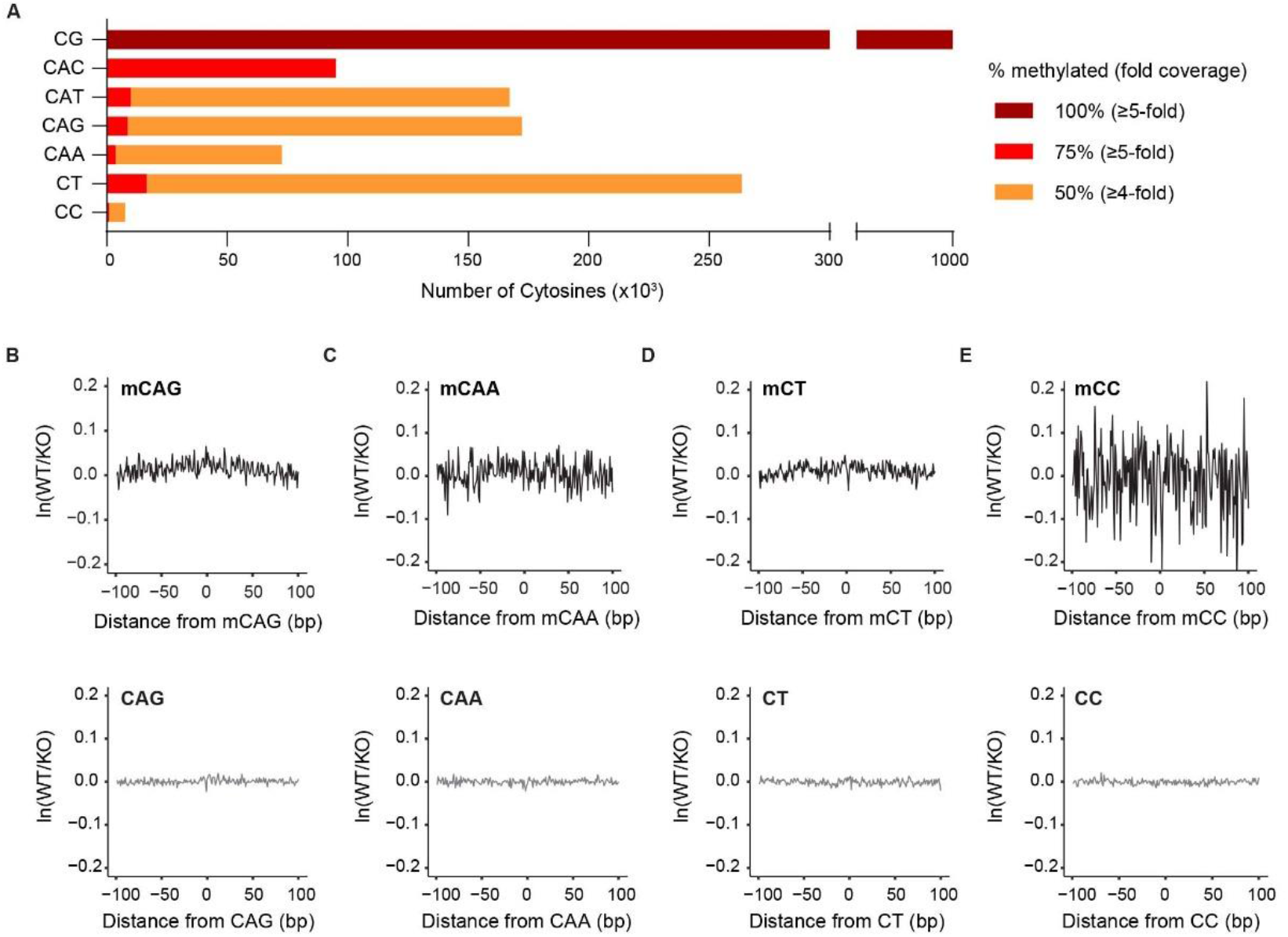
MeCP2 binds mCG and mCAC, and to a lesser extent, mCAT. A. Bar graph showing the number of highly methylated sites used for ATAC-seq footprinting. mCG sites are 100% methylated on both strands, with a combined ≥5-fold coverage. mCAC sites are ≥75% methylated, with a ≥5-fold coverage. All others are ≥50% methylated, with a ≥4-fold coverage. B-E. ATAC-seq footprints over methylated (upper) and unmethylated (lower) sequences. mCAG (B), mCAA (C), mCT (D) and mCC (E) footprints used sites with ≥50% methylation and ≥4-fold coverage, as shown in panel. All CH footprints used 1 million sites with 0% methylation and ≥10-fold coverage.

**Figure S2.**
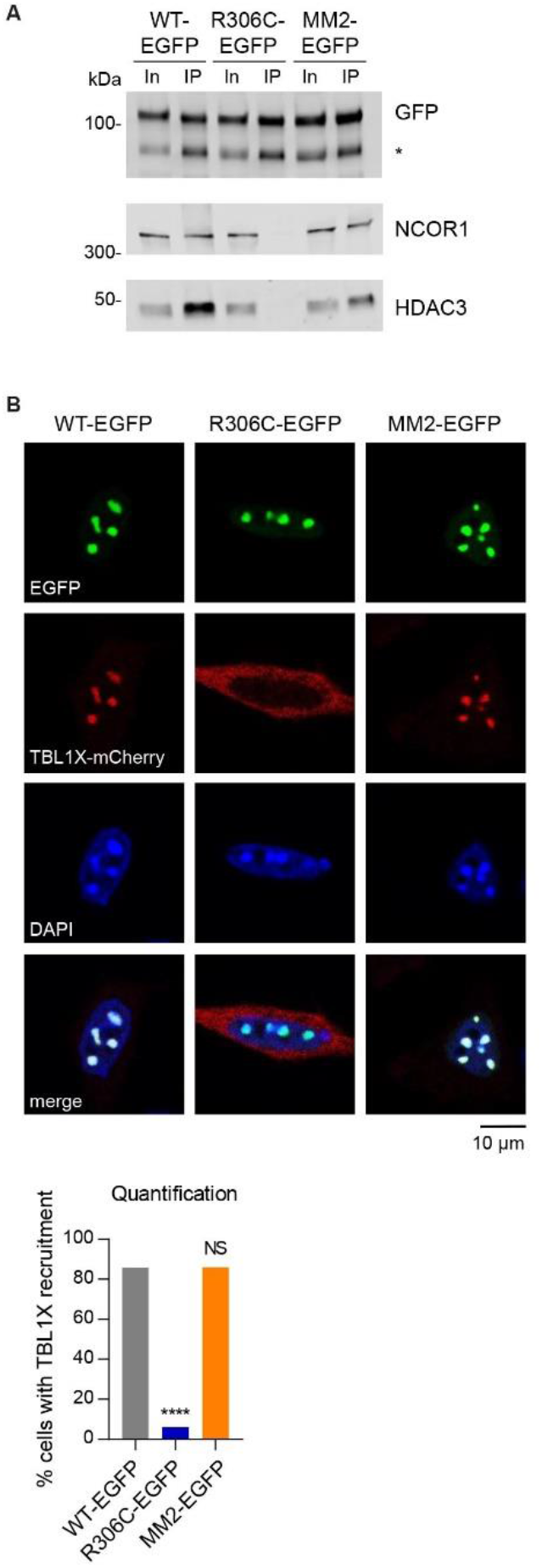
Chimeric protein MM2 retains the ability to bind to the key interaction partner of MeCP2, the NCOR1/2 repressor complexes. A. EGFP-tagged MM2 immunoprecipitated components of the NCOR1/2 co-repressor complexes: NCOR1 and HDAC3. WT and R306C were used as positive and negative controls for binding, respectively. In = input; IP = immunoprecipitate; * band from degraded protein. B. Representative images showing recruitment of TBL1X–mCherry to heterochromatic foci by EGFP-tagged MM2 when it is co-overexpressed in NIH-3T3 cells. WT and R306C were used as positive and negative controls for TBL1X–mCherry recruitment, respectively. Scale bar, 10 μm. Quantification (below) shows the percentage of cells with focal TBL1X–mCherry localization, evaluated relative to WT-EGFP using Fisher’s exact tests: R306C-EGFP, **** P < 0.0001; MM2-EGFP, P > 0.99. Total numbers of cells counted: WT-EGFP, n = 156; R306C-EGFP, n = 150; MM2-EGFP, n = 151; over three independent transfection experiments.

**Figure S3.**
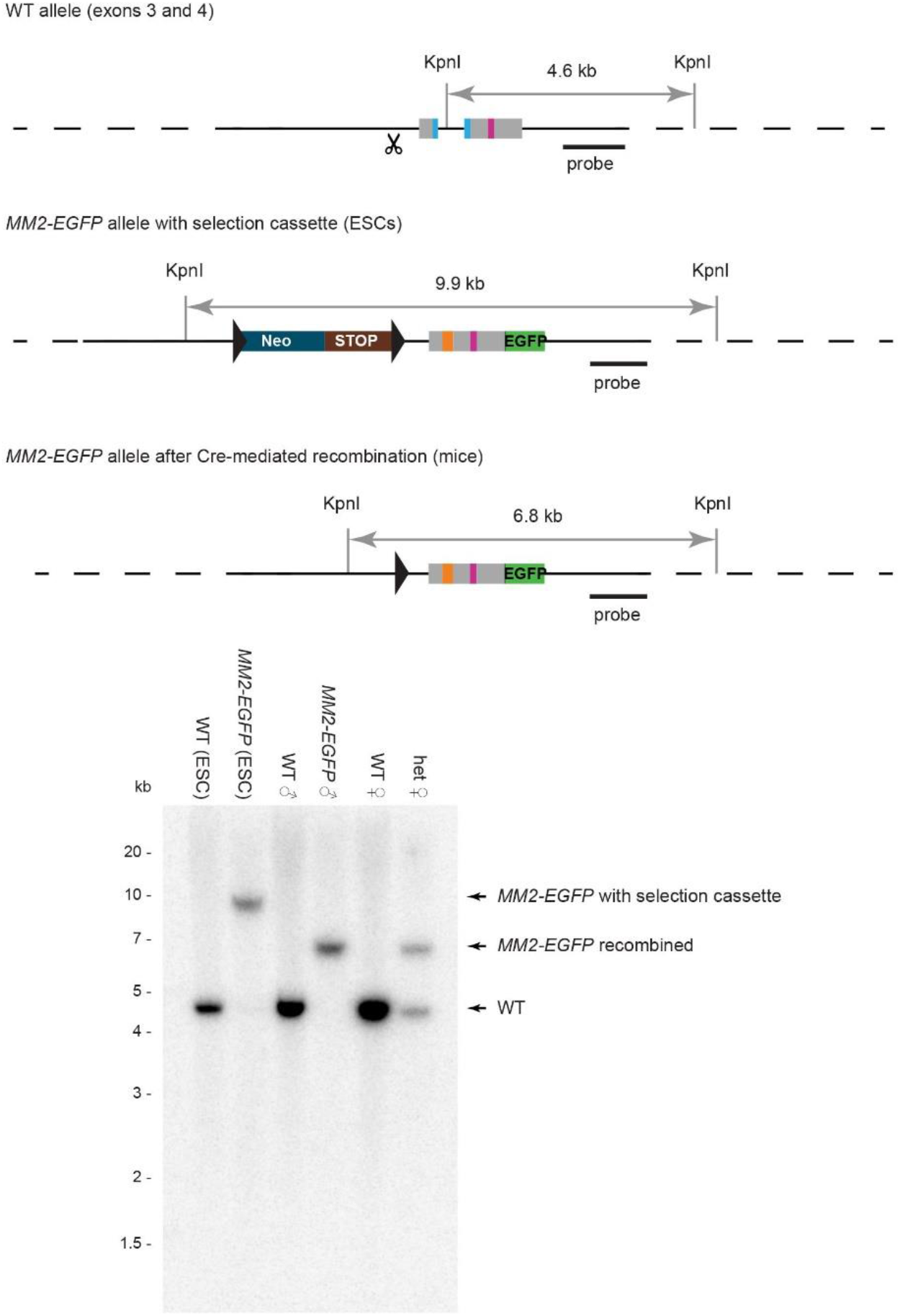
Generation of *MM2-EGFP* mice. Diagrammatic representation of *MM2-EGFP* mouseline generation. The endogenous *Mecp2* allele was targeted in male ES cells. The site of Cas9 cleavage in the WT sequence is shown by the scissors symbol. The floxed selection cassette was removed *in vivo* by crossing chimeras with deleter (*CMV-cre*) transgenic mice to produce constitutively expressing *MM2-EGFP* mice. The solid black line represents the sequence encoded in the targeting vector and the dotted lines indicate the flanking regions of mouse genomic DNA. *LoxP* sites are shown as triangles. Key: *Mecp2* MBD = blue, *MM2* MBD = orange, NID = pink, intervening *Mecp2* exonic sequences = grey, EGFP = green, linker = dark green, Neomycin resistance gene = dark blue, transcriptional stop cassette = dark red. Southern blot analysis shows correct targeting of ES cells and successful cassette deletion in the *MM2-EGFP* knock-in mice.

**Figure S4:**
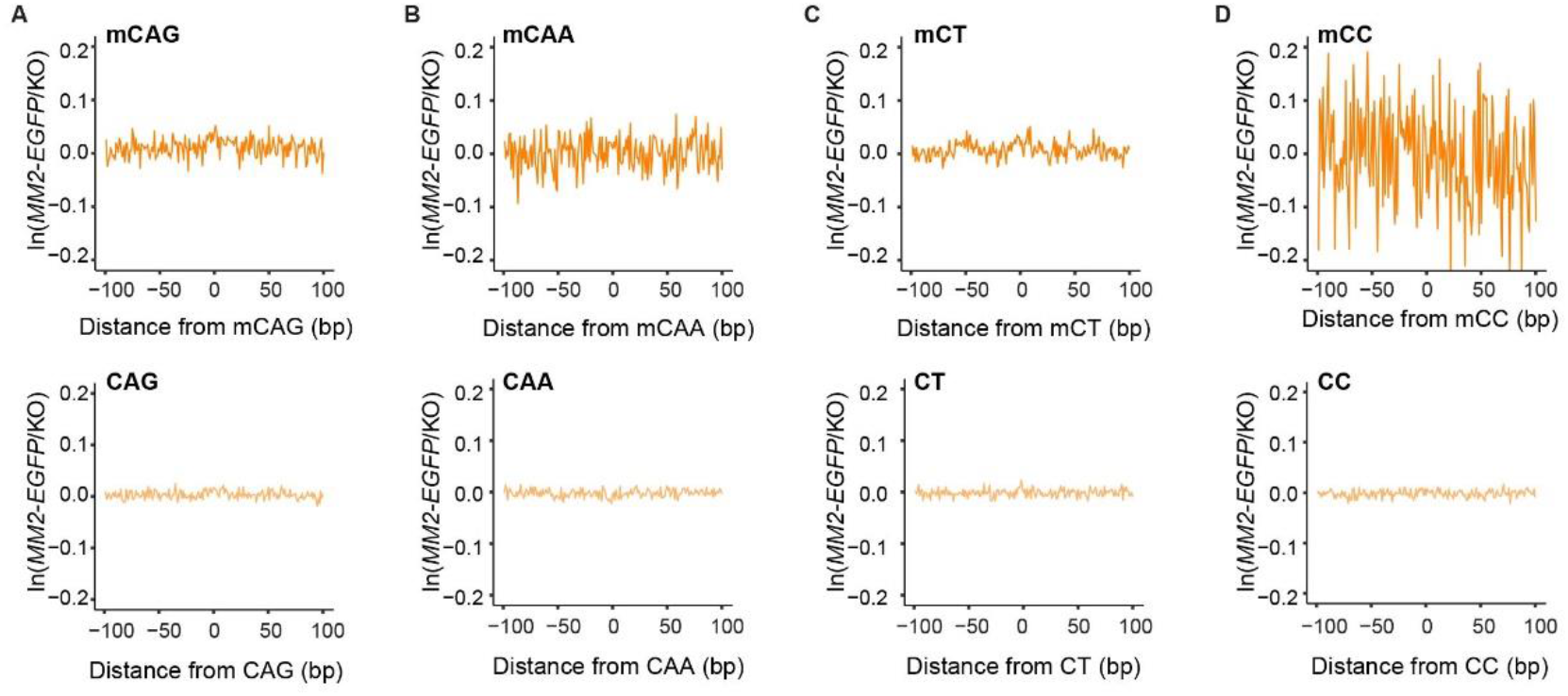
MM2 binds specifically to mCG *in vivo*. A-D. ATAC-seq footprinting of MM2-EGFP at methylated (upper) and unmethylated (lower) CAG (A), CAA (B), CT (C) and CC (D) sites. Equivalent WT ATAC-seq footprinting is shown in Figure S1B-E.

**Figure S5.**
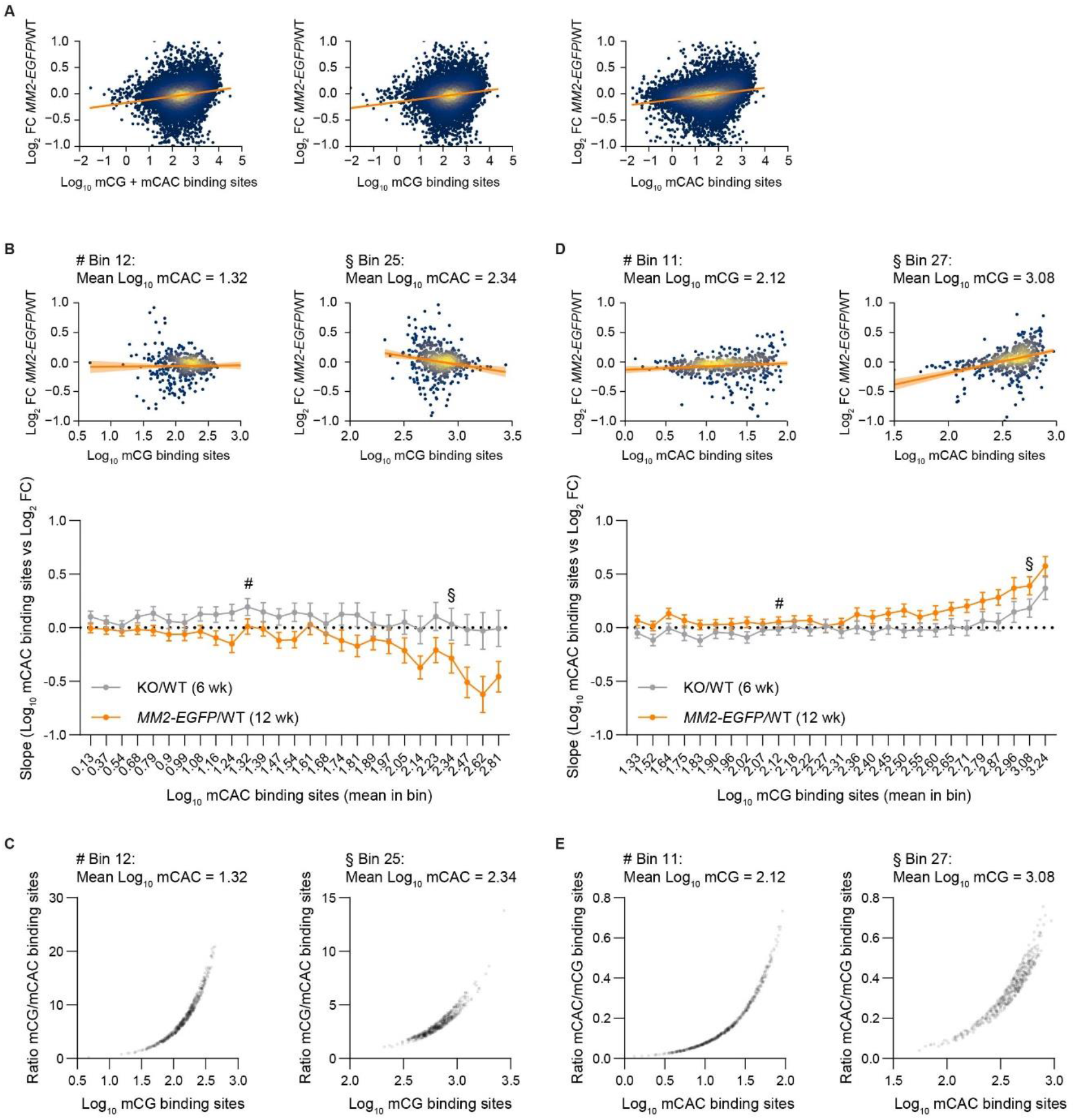
MM2 represses transcription at mCG but not mCAC sites. A. Correlations between the total number of MeCP2 binding sites per gene: mCG + mCAC (left), mCG (centre) and mCAC (right) and transcriptional changes in *MM2-EGFP*/WT hypothalamus tissue at 12 weeks of age. B. Genes were binned by the number of mCAC binding sites to determine the effect of mCG on transcription in *MM2-EGFP-EGFP/WT* (12 weeks). Two example bins are shown (# Bin 12, mean log_10_ mCAC = 1.32; and § Bin 25, mean log_10_ mCAC = 2.34). The slopes of all bins are shown below. C. The ratio of mCG/mCAC binding sites per gene in each mCAC bin. D. Genes were binned by the number of mCG binding sites to determine the effect of mCAC on transcription. Two example bins are shown (# Bin 11, mean log_10_ mCG = 2.12; and § Bin 27, mean log_10_ mCAC = 3.08). The slopes of all bins are shown below. Error on all slopes: 95% CI. E. The ratio of mCAC/mCG binding sites per gene in each mCG bin.

**Figure S6.**
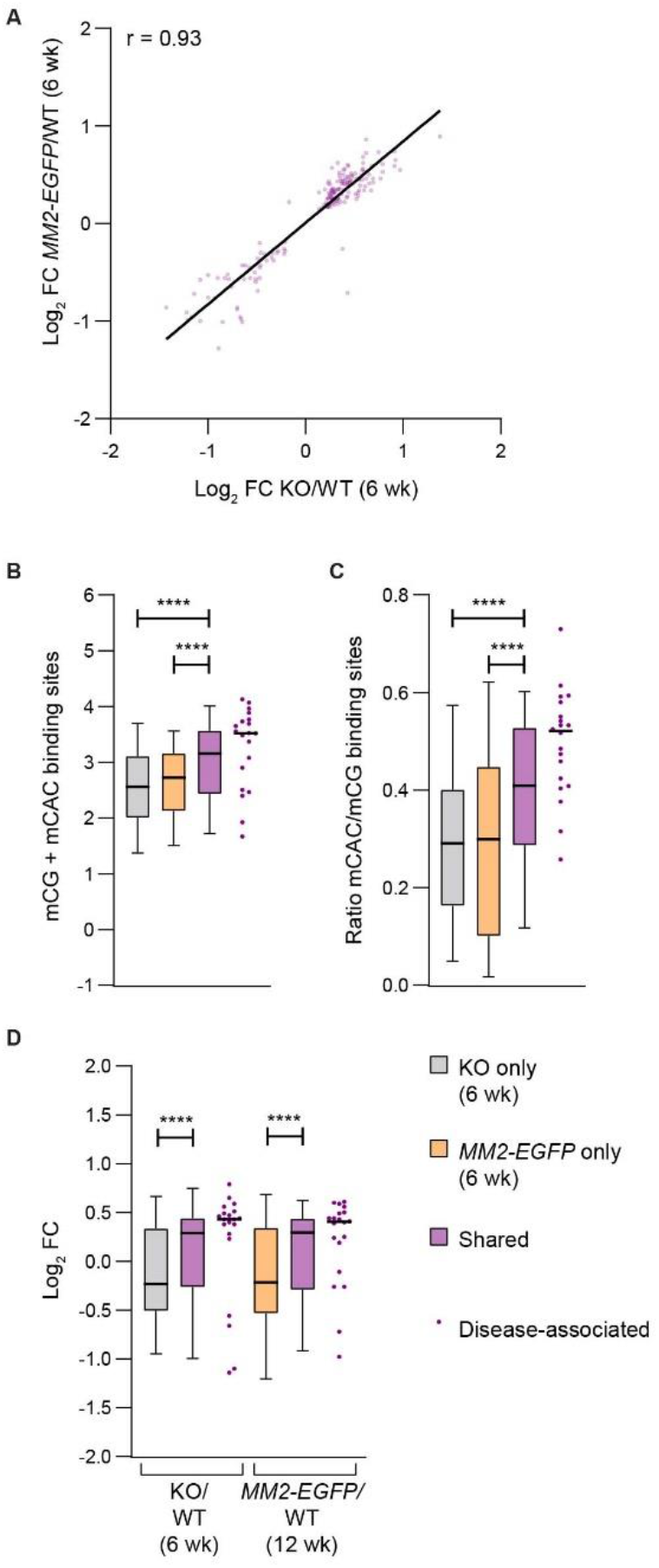
Shared dysregulated genes are enriched in mCAC and highly upregulated. A. Shared genes are dysregulated in the same direction in KO/WT (6 weeks) and *MM2-EGFP*/WT (6 weeks). B. Shared genes have more mCG + mCAC binding sites, compared to genes dysregulated in only KO or only *MM2-EGFP* (6 weeks). C. Shared genes have higher mCAC/mCG ratios, compared to genes dysregulated in only KO or only *MM2-EGFP* (6 weeks). D. Shared genes are more upregulated in both mutants, compared to genes dysregulated in only KO or only *MM2-EGFP* (6 weeks). B-D. Disease-associated genes have exceptionally high mCG + mCAC levels, mCAC/mCG ratios and are highly upregulated. Median values are indicated by a black bar. Pairs of gene sets were compared using Mann-Whitney tests: **** P < 0.0001.

**Table S1:** Disease Ontology analysis of shared dysregulated genes.

| DOID | Description | Gene ratio | BgRatio | P value | Adj. P | q value | Genes |
| --- | --- | --- | --- | --- | --- | --- | --- |
| 0060040 | pervasive developmental disorder | 16/155 | 201/8007 | 1.5E-06 | 0.00056 | 0.0005 | *MEF2C; RBFOX1; AVPR1A; SEMA5A; GRM8; AUTS2; GRIP1; DOCK4; NOS1AP; CNTNAP2; BDNF; OXTR; GRIN2A; NTNG1; TAC1; CNTN4* |
| 0060041 | autism spectrum disorder | 15/155 | 190/8007 | 3.6E-06 | 0.00056 | 0.0005 | *MEF2C; RBFOX1; AVPR1A; SEMA5A; GRM8; AUTS2; GRIP1; DOCK4; NOS1AP; CNTNAP2; BDNF; OXTR; GRIN2A; TAC1; CNTN4* |
| 12849 | autistic disorder | 15/155 | 190/8007 | 3.6E-06 | 0.00056 | 0.0005 | *MEF2C; RBFOX1; AVPR1A; SEMA5A; GRM8; AUTS2; GRIP1; DOCK4; NOS1AP; CNTNAP2; BDNF; OXTR; GRIN2A; TAC1; CNTN4* |
| 0060037 | developmental disorder of mental health | 20/155 | 373/8007 | 3.1E-05 | 0.00361 | 0.0035 | *MEF2C; RBFOX1; AVPR1A; SEMA5A; GRM8; AUTS2; KIRREL3; GRIP1; DOCK4; KCNK9; NOS1AP; CNTNAP2; CCSER1; CA8; BDNF; OXTR; GRIN2A; NTNG1; TAC1; CNTN4* |

## Key

DOID: Disease Ontology Unique Identification Number
Gene ratio: Ratio of genes which are part of the disease ontology in the query/input list (subset of the overlap of KO vs WT and MM2 vs WT which are associated with any disease term, not every gene is associated with disease ontology)
BgRatio: Ratio of genes which are part of the disease ontology in the background list (subset of all the genes that are associated with any disease ontology, not every gene is associated with disease ontology)
P-value: p-value associated with the hypergeometric test
Adj. P: Adjusted p-value after performing Benjamini-Hochberg correction
q value/FDR: False Discovery Rate after performing the Benjamini-Hochberg correction

## STAR METHODS

### CONTACT FOR REAGENT AND RESOURCE SHARING

Further information and requests for resources and reagents should be directed to and will be fulfilled by the Lead Contact, Adrian Bird.

All unique/stable reagents generated in this study are available from the Lead Contact without restriction.

The datasets generated during this study are available on GeoServer, accession number: GSE152801. The code for the analysis of fluorescence recovery after photobleaching (FRAP) data is available on GitHub DOI:10.5281/Zenodo.2654602

### EXPERIMENTAL MODEL AND SUBJECT DETAILS

#### Bacterial strains

BL21(DE3)RIPL competent cells (Agilent) were used to express MeCP2[MBD]. BL21(DE3)-R3-pRARE2 competent cells (Savitsky et al., 2010) were used to express MBD2[MBD].

BL21(DE3)pLysS competent cells (Promega) were used to express MeCP2[N-term] and MM2[N-term].

#### Cell lines

HeLa and NIH-3T3 (ATCC, RRID:CVCL_0594) cells were grown in DMEM (Gibco) supplemented with 10% fetal bovine serum (FBS; Gibco) and 1% penicillin– streptomycin (Gibco). Low passage male E14 TG2a ES cells (derived from 129/Ola mice) were a gift from Andrew Smith at the Centre for Genome Research at the University of Edinburgh (now Institute for Stem Cell Research). ES cells were grown in Glasgow MEM (Gibco) supplemented with 10% FBS (Gibco, batch tested), 1% non-essential amino acids (Gibco), 1% sodium pyruvate (Gibco), 0.1% β-mercaptoethanol (Gibco) and 1,000 units/ml LIF (ESGRO). Mouse ES cell status was confirmed regularly by production of germline chimeric mice from the cells (e.g. Tillotson et al., 2017). All cells were grown at 37°C, 5% CO_2_.

#### Mouse lines

All mice used in this study were bred and maintained at the Universities of Edinburgh animal facilities under standard conditions, and procedures were carried out by staff licensed by the UK Home Office and in accordance with the Animal and Scientific Procedures Act 1986. All mice were housed in a specific-pathogen-free (SPF) facility. They were maintained on a 12-h light/dark cycle and given ad libitum access to food (RM1 or RM3) and water. They were housed in open top cages with wood chippings, tissue bedding and additional environmental enrichment in groups of up to ten animals. Mutant mice were caged with their wild-type littermates.

*MM2-EGFP* mice were generated in this study from edited ES cells (detailed method described below). The edited allele contained a floxed selection cassette, which was removed by breeding chimeras with CMV-Cre deleter mice (C57BL/6J-Tg(CMV-cre)1Cgn, RRID: IMSR_JAX:006054). This line was further backcrossed onto C57BL/6J to N4 before behavioural phenotyping and to at least N4 for biochemical analysis. Hemizygous *MM2-EGFP* males (and wild-type controls) were used in all experiments as symptoms develop earlier and progress more rapidly than heterozygous females (as with other *Mecp2* mutant mice). To produce behavioural cohorts, male pups from timed matings were pooled at weaning and randomly assigned to the two groups. The first cohort was monitored for overt RTT-like symptoms, weight and survival from four weeks of age until each animal’s humane end-point (wild-type controls were monitored to one year of age). The second cohort underwent behavioural testing at 10-11 weeks of age. Tissues for biochemical analysis were harvested from animals at 6-13 weeks of age.

WT-EGFP (C57BL/6J-Mecp2^tm3.1Bird^, RRID: IMSR_JAX:014610) mice (Brown et al., 2016) were used as age- and sex-matched controls in some biochemical assays. They were maintained on a C57BL/6J background.

*Mecp2-null* mice (C57BL/6J;CBA/CA-Mecp2^tm2Bird^, RRID: IMSR_JAX:006849) mice (Guy et al., 2001) were used for ATAC-seq analysis to determine the footprints of wild-type MeCP2 and MM2-EGFP and for RNA-seq analysis. They were maintained on a mixed C57BL/6J;CBA/CA background, as breeding difficulties occur after backcrossing onto a C57BL/6J background. Hypothalamus tissue was harvested at 6 weeks of age.

Phenotypic data was compared to RTT models: *Mecp2-null* mice (C57BL/6J-Mecp2^tm2Bird^, RRID: IMSR_JAX:006849), *T158M-EGFP* (C57BL/6J-Mecp2^tm4.1Bird^, RRID:IMSR_JAX:026762), *R306C-EGFP* (C57BL/6J-Mecp2^tm5.1Bird^, RRID:IMSR_JAX:026847) and *R133C-EGFP* (C57BL/6J-Mecp2^tm6.1Bird^, RRID:IMSR_JAX:026848). This data was previously published in (Brown et al., 2016).

### METHOD DETAILS

#### Design of MM2

According to convention, all amino-acid numbers given refer to the e2 isoform. Numbers refer to homologous amino acids in human (NCBI accession P51608) and mouse (NCBI accession Q9Z2D6) protein, until residue 385 where there is a two-amino-acid insertion in the human protein. In this study, the MBD of mouse MeCP2 was defined as residues 94-164. This region was replaced by the MBD of mouse MBD2 (NCBI accession Q9Z2E1), comprising residues 153-217. Note: the region of MeCP2 replaced was shorter than the minimal fragment required for binding to methylated DNA (Nan et al., 1993). This region comprises the highly conserved sequence among MBD family members and contains all RTT-causing missense mutations in the MBD cluster, listed on RettBASE: http://mecp2.chw.edu.au/ (Krishnaraj et al., 2017). The region is flanked by proline residues (disrupting protein structure), which were selected as the junctions in MM2 (P93 and P165 of MeCP2). This protein was tagged at the C-terminus by EGFP, connected by a short linker. To be consistent with the WT-EGFP mice (Brown et al., 2016), the linker sequence was CKDPPVAT (DNA sequence: TGTAAGGATCCACCGGTCGCCACC). Intron 3 (located in the MBD of MeCP2) was not inserted into MM2 as it was deemed dispensable due to poor evolutionary conservation.

#### Plasmids

For expression of MeCP2[MBD] in bacteria, cDNA encoding MeCP2 (residues 77-167) was cloned into a pET28 using NdeI and EcoRI restriction sites. The protein sequence of this fragment is identical in human and mouse.

For expression of MBD2[MBD] in bacteria, cDNA encoding human MBD2 (residues 139-224, National Center for Biotechnology Information (NCBI) accession number NP_003918.1, from the Mammalian Gene Collection: BC032638.1, IMAGE: 5496721) was amplified by polymerase chain reaction (PCR) (primer sequences in Table S2) in the presence of Herculase II fusion DNA polymerase (Agilent Technologies). PCR products were purified (QIAquick PCR Purification Kit, QIAGEN UK) and further sub-cloned into a pET28-derived expression vector, pNIC28-Bsa4 using ligation-independent cloning (Savitsky et al., 2010). This vector includes sites for ligation-independent cloning and a Tobacco Etch Virus (TEV)-cleavable N-terminal His6-tag (extension MHHHHHHSSGVDLGTENLYFQ*SM-). After digestion with TEV protease, the protein retains an additional serine and methionine on the N terminus. The construct was transformed into competent Mach1 cells (Invitrogen, UK) to yield the final plasmid DNA.

For expression of MeCP2 WT and derivatives in mammalian cells, plasmids expressing EGFP-tagged e2 isoforms were used. Previously published (Tillotson et al., 2017) plasmids were used to express WT (RRID: Addgene_110186), R111G (RRID: Addgene_110187) and R306C (RRID: Addgene_110188). To clone MM2, the MBD of MBD2 and flanking MeCP2 sequences was synthesised (GeneStrings, Thermo Fisher Scientific; sequence in Key Resources table) and cloned into the pEGFP-N1_MeCP2[WT] plasmid to replace the MeCP2 MBD sequence using PstI restriction sites (NEB). R169G was inserted into the MM2 vector using a QuikChange II XL Site-Directed Mutagenesis Kit (Agilent Technologies) (primer sequences in Table S2). pmCherry-N1_TBL1X (Lyst et al., 2013) was used in the recruitment assay. For expression of MeCP2[N-term] and MM2[N-term] in bacteria for protein production, residues 1-205 of MeCP2 and the equivalent fragment of MM2 were subcloned from the pEGFP-N1 vectors into pET30b (Novagen). The forward primer introduced a NdeI restriction site and the reverse primer introduced a C-terminal 6xHis tag and EcoRI site (primer sequences in Table S2). To overcome low efficiency, the fragments were blunt cloned into pCR4 using the Zero Blunt TOPO kit (Invitrogen) before cloning into the pET30b vector using NdeI and EcoRI (NEB).

For targeting the endogenous *Mecp2* locus in ES cells, a gene-targeting construct containing the exons 3 and 4 of wild-type MeCP2 with a C-terminal tag (Tillotson et al., 2017) was edited by recombineering. The MBD region was first replaced by the counter-selection rpsL-neo cassette amplified by PCR to give 50 nt homology arms (primer sequences in Table S2). Positive clones were selected by kanamycin resistance. This cassette was subsequently replaced by the MBD of MBD2 flanked by MeCP2 sequences amplified from pEGFP-N1_MM2 (primer sequences in Table S2). Positive clones were selected by Streptomycin resistance and sequence verified. CRISPR– Cas9 technology was used to increase the targeting efficiency: the guide RNA sequence (GGTTGTGACCCGCCATGGAT) was cloned into pX330-U6-Chimeric_BB-CBh-hSpCas9 (a gift from F. Zhang; Addgene plasmid 42230 (Cong et al., 2013)).

All plasmids were sequence verified.

#### Generation of *MM2-EGFP* mice

The *MM2-EGFP* targeting vector was linearised using NotI (NEB). 129/Ola E14 TG2a ES cells were passaged every two days for one week and 20×10^6^ cells were collected in 600 μl HBS containing 15 μg linearised targeting vector and 15 μg of the CRISPR/Cas9 plasmid. The cells were electroporated (GenePulser X, Biorad) in a 0.4 cm cuvette at 240 V, 500 μF, ∞ resistance. The guide RNA targeted intron 2 of the wild-type gene (at the site of the NeoSTOP cassette in the targeting vector). G418-resistant clones with correct targeting at the *Mecp2* locus were identified by PCR and Southern blot screening (method described below). PCR screens were performed using Phusion polymerase (NEB): primer sequences in Table S2. Screening identified one positive clone (5C7), which was further verified by Sanger sequencing of the *Mecp2* locus and the top predicted intragenic off-target locus, *Dock5* (primer sequences in Table S2). The cell line was also karyotyped by inducing metaphase arrest by treatment for 3 hours with 0.1 μg/ml colemid (KARYOMAX, Gibco). Pelleted cells were resuspended in 300 μl growth media plus 5 ml 0.4% (w/v) KCl and incubated for 10 min. 100 μl fixative (3 parts methanol: 1 part acetic acid) was added and mixed gently. Cells were pellets at 300 *g*, 5 min and the pellet was resuspended in 5 ml fixative and incubated at room temperature for 20 min. Cells were pelleted again and resuspended in 500 μl fixative. They were dropped onto pre-chilled glass slides from ~30 cm and left to dry overnight before mounting with Vectashield containing DAPI (H-1200). Cells were photographed on a Zeiss AxioImager (Carl Zeiss UK, Cambridge) equipped with a Photometrics Prime sCMOS camera (Teledyne Photometrics) and chromosomes were counted manually using ImageJ (NIH).

Cells from *MM2-EGFP* clone 5C7 were injected into blastocysts (E3.5) obtained from C57BL/6J females after natural mating. As with previous studies, e.g. (Guy et al., 2001), 15 ES cells were injected into each blastocyst and 12 injected blastocysts were transferred to pseudo-pregnant recipient females (F1 C57BL/6J;CBA/CA). Chimeric pups were recognised by coat colour. This targeted locus contained a neomycin resistance gene followed by a transcriptional STOP cassette flanked by *loxP* sites (‘floxed’) in intron 2. This cassette was removed *in vivo* by crossing chimaeras with homozygous females from the transgenic CMV-cre deleter strain (Tg(CMV-cre)1Cgn, RRID: IMSR_JAX:006054) on a C57BL/6J background. The CMV-cre transgene was subsequently bred out. Genotyping primers for the Mecp2 locus and Cre transgenes are in Table S2.

#### Southern blotting

Genomic DNA was purified from ES cells using Puregene Core Kit A (Qiagen) according to manufacturer’s instructions. Genomic DNA was purified from snap-frozen mouse brain (harvested at 7-19 weeks of age) by homogenisation in 3 ml lysis buffer (50 mM Tris HCl pH7.5, 100 mM NaCl, 5 mM EDTA). Proteinase K was added to a final concentration of 0.4 mg/ml and SDS to 1%. After overnight incubation at 55°C, samples were treated with 0.1 mg/ml RNase at 37°C for 1-2 hours and gDNA was extracted with 3 ml PCI (phenol:chloroform:isoamyl alcohol, Sigma). gDNA was precipitated from the aqueous phase with 2.5 volumes 100% ethanol and 0.1 volumes 3M NaOH, and dissolved in TE. For Southern blotting, 10 μg gDNA was digested with 3 μl enzyme (NEB) in a total volume of 40 μl, and ran on 0.8% gels. Gels were incubated in 0.25 M HCl (15 min) and neutralised with 0.4 M NaOH (45 min) before transferring onto ZetaProbe membranes (Biorad). 25 ng of a probe homologous to the 3’ homology arm (1038 pb fragment digested with SpeI and BamHI) was radioactively labelled with [α^32^]dCTP (Perkin Elmer) using the Prime-a-Gene Labelling System (Promega) according to manufacturer’s instructions. Membranes with blocked in Church buffer containing ~50 μg/ml Herring Sperm DNA (Sigma) at 65°C. After 30 min, the probe was added and incubated overnight. Membranes were then washed three times with 3xSSC/0.2% SDS, followed by up to three washes with 1xSSC/0.2% SDS. The blots were exposed for 1-5 days on Phosphorimager plates (GE Healthcare) and scanned using a Typhoon FLA 7000. BamHI, BsrGI and KpnI were used to screen ES cell clones and KpnI was used to verify deletion of the NeoSTOP cassette in *MM2-EGFP* knock-in mice

#### Protein purification

The expression plasmid containing MeCP2[MBD] was transformed into competent BL21 (DE3)-RIPL cells (Life Technologies) and the MBD2[MBD] plasmid was transformed into competent BL21 (DE3)-R3-pRARE2 cells (a phage-resistant derivative of the BL21 (DE3) strain), with a pRARE plasmid encoding rare codon tRNAs (Invitrogen). Freshly grown colonies were cultured overnight in lysogeny broth (LB) supplemented with 50 mg/mL kanamycin and 34 mg/mL chloramphenicol at 37°C. One litre of pre-warmed terrific broth (TB) was inoculated with 10 mL of the overnight culture and incubated at 37°C. At an optical density at 600 nm (OD600) of 2.5, the culture was cooled to 18°C and expression was induced overnight at 18°C with 0.1 mM isopropyl-b-D-thiogalactopyranoside (IPTG). Cells were harvested by centrifugation (8700 x g, 15 min, 4°C) in a Beckman Coulter Avanti J-20 XP centrifuge, and then re-suspended in lysis buffer (50 mM HEPES pH 7.5 at 20°C, 1500 mM NaCl, 5% Glycerol, 1 mM tris(2-carboxyethyl)phosphine (TCEP) and 1:1000 (v/v) Protease Inhibitor Cocktail III (Calbiochem)). Cells were lysed using a high-pressure homogeniser (Emulsiflex C5, Avestin) at ~100 mPa equipped with a recirculating cooler (F250, Julabo). DNA was precipitated on ice for 30min with 0.15% (v/v) PEI (Polyethyleneimine, Sigma) and the lysate was cleared by centrifugation (16,000 x g for 1 hour at 4C, JA 25.50 rotor, Beckman Coulter Avanti J-20 XP centrifuge) before being applied to a Nickel affinity column (nickel nitrilotriacetic acid (Ni-NTA) resin, QIAGEN Ltd., 5 ml, equilibrated with 20 ml lysis buffer). Protein was eluted with an imidazole step gradient (50 to 250 mM) in a 500 mM NaCl lysis buffer.

For MeCP2[MBD], the eluted protein after Ni-NTA purification was concentrated with 10kDa MWCO Amicon® Ultra (EMD Millipore) concentrators and further purified by size exclusion chromatography on a HiPrep 16/60 Sephacryl S-200 gel filtration column (GE Healthcare) on an ÄktaPrime plus systems (GE/Amersham Biosciences). For MBD2[MBD], the eluted protein after Ni-NTA affinity purification was treated overnight at 4°C with TEV protease to remove the 6×His tag and untagged proteins were further purified by size exclusion chromatography on a Superdex 75 16/60 HiLoad gel filtration column (GE Healthcare Life Sciences). All eluted fractions were monitored by SDS-PAGE and concentrated in gel filtration buffer (10 mM HEPES pH 7.5, 500 mM NaCl and 5% glycerol) using Amicon® Ultra (EMD Millipore) concentrators with a 10 kDa MWCO cut-off. Proteins were aliquoted, flash frozen in liquid nitrogen and stored at −80°C until further use.

Expression plasmids containing MeCP2[N-term] and MM2[N-term] with a C-terminal 6xHis tag were transformed into BL21(DE3)pLysS competent cells and plated. Colonies were scraped into an overnight starter culture in LB with 50 μg/ml kanamycin and 17 μg/ml chloramphenicol at 37°C. This was expanded in 500 ml and protein production with induced with 1 mM IPTG at 30°C for 3 hours when the OD_600_ reached 0.6-0.8. Frozen bacterial cell pellets were mashed in 30 ml ice-cold lysis buffer (50 mM NaH_2_PO_4_, 100 mM NaCl, 10% glycerol, 30 mM imidazole, 0.1% NP40, and protease inhibitor tablet (Roche), 5.7 mM β-mercaptoethanol) and passed through a 21G needle. 750 U benzonase (Sigma) was added samples were sonicated for 10 cycles of 30 s ON/OFF at 30% amplitude (Branson Digital Sonifier). NaCl concentration was increased to 300 mM and samples were centrifuged at 31,000 *g* (30 min, 4°C) and the supernatant was transferred to new tubes. His-tagged proteins were purified using 0.5 ml NiSO_4_-coated Chelating Sepharose Fast Flow beads (GE healthcare), incubated in lysates for 1 hr at 4°C. Beads were washed three times in 12 ml lysis buffer (with 300 mM NaCl) and protein was eluted in five fractions each of 0.5 ml lysis buffer with 250 mM imidazole. Fractions were pooled and diluted with 5-10 ml HEPES buffer (20 mM HEPES, 300 mM NaCl, 1 mM EDTA). The samples subsequently purified using a HiTRap Sp Hp 1 ml Column (GE Healthcare) to select for positively charged proteins. Columns were washed with 10 ml HEPES buffer and protein was eluted in 1 ml HEPES buffer with 0.7 M NaCl (fraction 1) and then 1 ml 1 M NaCl (fraction 2). Fraction 2 was used in the EMSA experiments.

#### Bio-Layer Interferometry (BLI)

Single-stranded DNA probes were purchased from Integrated DNA Technologies (IDT) and annealed by heating at 95°C for 5min and slowly cooled down at room temperature. See Key Resources Table for sequences.

The dissociation constant (K_D_) of MeCP2[MBD] to different DNA probes was determined by using bio-layer interferometry on the Octet RED384 system (ForteBio, Pall). All assays were performed in low-binding black 96 well plates (Greiner) with 1,000 rpm orbital shaking and samples were diluted in freshly prepared and filtered BLI buffer (150mM NaCl, 10mM HEPES, pH8.0 and 0.05% Tween20).

First, streptavidin biosensors (18-5019, ForteBio, Pall) were hydrated in 200uL of BLI buffer for 20min at room temperature. Then, 5’-biotinylated double-stranded DNA probes (6nM, final concentration) were loaded on the streptavidin biosensors for 600sec followed by quenching with biotin (5ug/mL, final concentration) for 50sec. The DNA-loaded sensors were then submerged in wells containing increasing concentrations of MeCP2[MBD] (0-375nM) in the presence of poly(dI-dC) (1ug/mL, final concentration) (LightShift, Thermo) for 600sec followed by 300sec of dissociation time.

#### Electromobility shift assay (EMSA)

For the EMSA with MBD2[MBD2], single-stranded FAM-labelled DNA probes were purchased from Integrated DNA Technologies (IDT) and annealed by heating at 95°C for 5 min and slowly cooled down at room temperature. See Key Resources Table for sequences. Assays were assembled in 10 μL reactions in binding buffer containing 20 mM HEPES (pH7.9), 150mM KCl, 8% Ficoll, 1 mM EDTA and 0.5 mM DTT. The protein (2 μM final concentration) was incubated in binding buffer for 10min at room temperature, then poly(dI-dC) (LightShift, Thermo) was added at a final concentration of 40 ng/uL and the reaction incubated for 10 min at room temperature. 5’-FAM-labelled DNA probe (10 nM final concentration) was added and incubated a further 25 min at room temperature. The reactions were run on a 4.0% acrylamide (ratio 37:1, Sigma) Tris Borate EDTA gel in 0.25X TBE, 150V for 3 hours at 4°C. The bandshifts were exposed at 473 nm on a fluorescent scanner (FujiFilm FLA-5100) and gel images were visualized with the software FujiFilm MultiGauge.

EMSAs with MeCP2[N-term] and MM2[N-term] were performed as described previously (Brown et al., 2016). Single-strand oligonucleotide probes from the mouse *Bdnf* promoter region were purchased from Biomers, Germany, annealed by heating at 100 °C for 10 min and slowly cooled to room temperature. Probe sequences contained a central methylated, or unmethylated CG or CAC site (see Key Resources Table for sequences). 500 ng of probe was radio-labelled using T4 polynucleotide kinase (NEB) and purified (MinElute PCR Purification Kit, Qiagen) according to manufacturer’s instructions. 1 ng probe and 1 μg poly deoxyadenylic-thymidylic acid competitor DNA (Sigma-Aldrich) were added with 0, 1.5, 3.0 or 4.5 uM polypeptide in reaction buffer (5% glycerol, 0.1 mM EDTA, 10mM Tris HCl pH 7.5, 150 mM KCl, 0.1 mg/ml BSA) on ice for 20 minutes. Samples were run at 120 V for 70 min on a 10% acrylamide Tris Borate EDTA gel (0.075% APS, 0.00125% TEMED) in chilled TBE. The gels were exposed overnight and imaged using the Typhoon FLA 9500 scanner (GE Healthcare).

#### DNA pull-down assay

This assay performed as described previously (Connelly et al., 2020)(Piccolo et al., 2019). PCR-generated, biotin end-labelled 147 bp DNA probes (2 μg) were coupled to M280-streptavidin Dynabeads according to the manufacturer’s instructions (Invitrogen). All cytosines in the probes were either non-methylated or methylated and only occurred in a single sequence context (CG, CAC or CAT), sequences in the Key Resources table. Bead-DNA complexes were then co-incubated with 20 μg of rat brain nuclear protein extract (Mellén et al., 2017) for 1.5 hours at 4°C. Following extensive washing, bead-bound proteins were eluted using Laemmli buffer (Sigma) and resolved on a 4–15% SDS-polyacrylamide gel (NEB). The presence of MeCP2 was assayed by western blot using MeCP2 (Sigma M6818; RRID:AB_262075) diluted 1:1000; with secondary detection employing Li-COR IRDye 800CW Donkey anti-Mouse (926-32212) diluted 1:10,000, then scanned using a LI-COR Odyssey CLx machine.

#### MeCP2 localization and TBL1X–mCherry recruitment assay

Analysis of MeCP2 localisation and recruitment of TBL1X was performed as done previously (Tillotson et al., 2017). NIH-3T3 cells were seeded on coverslips in six-well plates (25,000 cells per well) and transfected with 2 μg plasmid DNA (pEGFP-N1–MeCP2 alone or pEGFP-N1–MeCP2 and pmCherry-N1–TBL1X5) using JetPEI (PolyPlus Transfection). After 48 h, cells were fixed with 4% (w/v in PBS) paraformaldehyde (Sigma), stained with 4′, 6-diamidino-2-phenylindole (DAPI; Sigma) and then mounted using ProLong Diamond (Life Technologies). Images were acquired on a Leica SP5 laser scanning confocal and a HCX PLAPO 63x/1.4 objective with laser lines for DAPI (405nm), GFP (488nm) and mCherry (594nm) selected using LAS AF software (Leica).

#### Fluorescence recovery after photobleaching (FRAP)

NIH-3T3 cells were seeded on poly-L-lysine-coated coverslips in six-well plates (200,000 cells per well) and transfected with 2 μg plasmid DNA using JetPEI (PolyPlus Transfection). After 40-48 h, coverslips were used for live cell imaging at 37°C, 5% CO2. Images were acquired on a Zeiss LSM880 laser scanning confocal equipped with a GaSP detector and a Plan-Apochromat 63x/1.4 objective. A z-stack was acquired at each time point comprising of six Z positions spaced 1 μm apart to account for any vertical movement of the bleach position in the cell. Images were acquired using a 488 nm laser at 0.2% power to minimise any photobleaching, except for the bleach spot (diameter 10 pixels = 1.6 μm) that was exposed to 100% laser power for 0.5 s (sufficient to produce an 80% decrease in the intensity at the bleach spot). A total of 295 post bleach images (1.02 s apart) were taken to ensure any recovery was measured to a steady state. Five images were taken before the bleach, the mean of which was used as the “preBleach” values. At least two cells were included in each frame, the first containing the bleach spot and the second as a control to correct for photobleaching during the course of the experiment.

#### Immunohistochemistry

Mice were culled using CO_2_, and as soon as death could be confirmed, the rib cage was opened up to expose the heart. Animals were perfused with ~1ml Lidocaine (1% in PBS) followed by ~25 ml 4% paraformaldehyde (w/v) in PBS (Sigma) pumped at 7 ml/min through the circulatory system. Brains were stored overnight in 4% PFA (w/v) in PBS before incubation in 30% sucrose (w/v) in PBS overnight and flash freezing in isopentane. 10 μm sections were cut on a Leica CM1900 cryostat at −18°C. Sections were washed twice in PBS, permeablised in 0.1% (v/v) Triton-100 in PBS (15 min), washed again and blocked in 1.5% goat serum. They were then probed with a Cy3-conjugated antibody against NeuN (Millipore MAB377C3; RRID: AB_10918200) overnight at 4°C. Sections were washed once in PBS, incubated in 1 μg/ml DAPI (10 min), washed twice more and mounted with coverslips using Prolong Diamond (ThermoFisher Scientific). Images of sections were acquired on a Leica SP5 laser scanning confocal and a HCX PLAPO 63x/1.4 objective with laser lines for DAPI (405nm), GFP (488nm) and Cy3 (561 nm) selected using LAS AF software (Leica). Images show neurons in the dentate gyrus of the hippocampus.

#### Immunoprecipitation and western blotting

HeLa cells were transfected with pEGFP-N1–MeCP2 plasmids using JetPEI (PolyPlus Transfection) and harvested after 24–48 h. Fresh or frozen cell pellets were Dounce homogenised in 1.5 ml in NE1 (20 mM HEPES pH 7.5; 10 mM NaCl; 1mM MgCl_2_; 0.1% Triton X-100; 10 mM β-mercaptoethanol; protease inhibitor tablet (Roche)). Nuclei were collected by centrifugation at 845 *g* (5 min, 4°C) and resuspended in 200 μl NE1. Nuclei were treated with 250 U Benzonase (Sigma) for 5 min at room temperature before increasing NaCl concentration to 150 mM and incubated under rotation for 20 min at 4°C. Samples were centrifuged at 16,000 *g* (20 min, 4°C) and the supernatant was taken as the nuclear extract. MeCP2–EGFP complexes were captured from nuclear extract using GFP-Trap_A beads (Chromotek) under rotation at 4°C for 20-30 min. Beads were washed four times in NE1 containing 150 mM NaCl and then proteins were eluted in 2x Laemmli Sample Buffer (Sigma) at 100°C, 5 min. Proteins were analysed by western blotting using antibodies against GFP (NEB 2956; RRID: AB_1196615), NCOR1 (Bethyl A301-146A; RRID: AB_873086) and HDAC3 (Sigma 3E11; RRID: AB_1841895), all at a dilution of 1:1,000; followed by LI-COR secondary antibodies: IRDye 800CW Donkey anti-Mouse (926-32212) and IRDye 800CW Donkey anti-Rabbit (926-32213) or IRDye 680LT Donkey anti-Rabbit (926-68023) at a dilution of 1:10,000. Signal was detected on a LI-COR Odyssey CLx machine.

Protein levels in whole-brain crude extracts were quantified using western blotting. Frozen half-brains (harvested at 9 weeks of age) were Dounce homogenised in 750 μl cold NE1 (20 mM HEPES pH7.9, 10 mM KCl, 1 mM MgCl2, 0.1% Triton X-100, 20% glycerol, 0.5 mM DTT, and protease inhibitors (Roche)) and treated with 750 U Benzonase (Sigma) for 15 min at room temperature. Next, 750 μl of 2x Laemmli Sample Buffer (Sigma) was added and samples were boiled for 3 min at 100°C, snap-frozen on dry ice, and boiled again for 5 min. Samples were diluted 1/6 with 1x sample buffer before analysis by western blotting. Membranes were probed with an antibody against MeCP2 (Sigma M7443; RRID:AB_477235) at a dilution of 1:1,000, followed by LI-COR secondary antibodies (listed above). Histone H3 (Abcam ab1791) was used as a loading control (dilution 1:10,000).

#### Flow cytometry

Fresh brains were harvested from 12-week-old animals and Dounce-homogenized in 5 ml homogenization buffer (320 mM sucrose, 5 mM CaCl2, 3 mM Mg(Ac)2, 10 mM Tris HCl pH 7.8, 0.1 mM EDTA, 0.1% NP40, 0.1 mM PMSF, 14.3 mM β-mercaptoethanol, protease inhibitors (Roche)), and 5 ml 50% OptiPrep gradient centrifugation medium (50% Optiprep (Sigma D1556-250ML), 5 mM CaCl2, 3 mM Mg(Ac)2, 10 mM Tris HCl pH 7.8, 0.1 M PMSF, 14.3 mM β-mercaptoethanol) was added. This was layered on top of 10 ml 29% OptiPrep solution (v/v in H2O, diluted from 60% stock) in Ultra-Clear Beckman Coulter centrifuge tubes, and samples were centrifuged at 10,100 *g* (30 min, 4 °C). Pelleted nuclei were resuspended in resuspension buffer (20% glycerol in DPBS (Gibco) with protease inhibitors (Roche)). For flow cytometry analysis, nuclei were pelleted at 600 *g* (5 min, 4 °C), washed in 1 ml PBTB (5% (w/v) BSA, 0.1% Triton X-100 in DPBS with protease inhibitors (Roche)) and then resuspended in 250 μl PBTB. To stain for NeuN, NeuN antibody (Millipore MAB377) was conjugated to Alexa Fluor 647 (APEX Antibody Labelling Kit, Invitrogen A10475), added at a dilution of 1:125 and incubated under rotation for 45 min at 4 °C. Flow cytometry (BD LSRFortessa SORP using FACSDIVA version 8.0.1 software) was used to obtain the mean EGFP fluorescence for the total nuclei (n = 50,000 per sample) and the high-NeuN (neuronal) subpopulation (n > 8,000 per sample). Three biological replicates of each genotype were analysed.

#### ChIP-qPCR

Snap-frozen brain hemispheres (harvested at 8-9 weeks of age) were homogenised for 30 s using a Polytron in 7 ml crosslinking buffer (2% formaldehyde, 10 mM HEPES pH 7.5, 100 mM NaCl, 1 mM EDTA, 1 mM EGTA) and incubated for 30 s at room temperature before quenching with 7 ml 350 mM glycine for 5 min on rocker. Samples were centrifuged at 2000 *g* (3 min, 4°C) and pellets were resuspended in 2 ml L1 buffer (50 mM HEPES pH 7.5, 140 mM NaCl, 1 mM EDTA, 1 mM EGTA, 0.25% Triton X-100, 0.5% NP-40, 10% glycerol, protease inhibitors (Roche)) and Dounce homogenised (tight pestle). Another 3 ml L1 buffer was added and samples were centrifuged at 2000 *g* (10 min, 4°C). Pellets were resuspended in 5 ml L1 buffer again, then centrifuged again. Pellets were resuspended in 5 ml L2 buffer (10 mM Tris HCl PH 8.0, 200 mM NaCl, protease inhibitors (Roche)) and rotated for 10 min at room temperature. Samples were centrifuged again and pellets were resuspended in 1 ml L3 buffer (10 mM Tris HCl PH 8.0, 1 mM EDTA, 1 mM EGTA, protease inhibitors (Roche)) with 0.1% SDS. Samples were sonicated in 300 μl aliquots for 22 cycles of 30 s ON/OFF on high power on a Bioruptor Sonicator (Diagenode). Lysates were cleared by centrifugation at 16,000 *g* (10 min, 4C°), and 30 μg chromatin was used per IP. This was topped up to 400 μl with L3 buffer and 200 μl 3X Covaris ChIP buffer (20 mM Tris HCl PH 8.0, 3% Triton X-100, 450 mM NaCl, 3 mM EDTA) was added. Lysates were precleared by incubation with 10 μl pre-washed blocked magnetic agarose beads (Chromotek) under rotation for 1 hour at 4°C. Inputs were taken at this stage (12 μl = 2%). 25 μl pre-washed GFP-Trap magnetic agarose beads (Chromotek) were added to immunoprecipitation EGFP-tagged proteins under rotation at 4°C overnight. Beads were washed 500 μl ice-cold low salt buffer (0.1% SDS, 1% Triton X-100, 20 mM Tris HCl pH 8.0, 150 mM NaCl, 2 mM EDTA) twice, high salt buffer (0.1% SDS, 1% Triton X-100, 20 mM Tris HCl pH 8.0, 500 mM NaCl, 2 mM EDTA) twice and TE. Chromatin was recovered by incubating beads twice in 100 μl elution buffer (1% SDS in TE) at 65°C for 30 min shaking at 1200 rpm. Inputs were made up to 200 μl with elution buffer. IPs and inputs were incubated at 65°C overnight to de-crosslink. Samples were treated with 50 μg/ml RNaseA for 1 hour at 37°C and then 0.7 mg/ml Proteinase K for 2-3 hours at 55°C. To purify DNA, 200 μl Phenol/Chloroform/Isoamylethanol (PCI) was added and samples were transferred to 5PRIME Phase Lock at centrifuged at 16,000 *g* (5 min, 4C°). DNA was extracted from the aqueous phase using the Qiagen PCR purification kit according to manufacturer’s instructions, eluted in 2X 30 μl water and diluted to 180 μl. qPCR was performed at three mCG-rich/mCA-poor loci: Cyp3a25, Cyp3a57 and Cyp3a59. The reaction mixture consisted of 2.5 μl DNA, 500 nM each primer, 1X SYBR Green I (Sigma) in a total volume of 12.5 μl. Primer sequences in Table S2 (Brown et al., 2016). Standard curves were obtained for each primer pair using a 2-fold dilution of DNA. Samples were processed on a LightCycler 480: 95°C 10 min; 45 cycles of 95°C 15 s, 58°C 15 s, 72°C 20 s with single acquisition mode).

#### ATAC-seq library preparation

Hypothalami were dissected from *MM2-EGFP* (n = 3), *Mecp2*-null (KO) (n = 3) and WT (n = 4, 3 *MM2*-EGFP littermates and 1 KO littermate) mice at 6-7 weeks of age. Freshly dissected tissues were homogenised using a hypotonic buffer (10 mM Tris-HCl pH 7.4, 10 mM NaCl, 3mM MgCl_2_, 0.1% [v/v] Igepal CA-630). Isolated nuclei were counted, and 50,000 nuclei were resuspended in 50 μl of a transposition reaction mix containing 2.5 μl Nextera Tn5 Transposase and 2x TD Nextera reaction buffer. The mix was incubated for 30 min at 37 °C. DNA was purified by either the MinElute PCR kit (Qiagen) or the Agencourt AMPure XP beads (Beckman Coulter) and PCR amplified with the NEBNext High Fidelity reaction mix (NEB) to generate DNA libraries. The libraries were sequenced as 75 bp paired-end reads on a HiSeq 2500 Illumina platform.

#### RNA-seq library preparation

Hypothalami were harvested at 6-7 weeks of age (referred to as “6 weeks”) from *Mecp2*-null (KO) mice (n = 4) and their WT littermate controls (n = 3) and *MM2-EGFP* mice (n = 3) and their WT littermate controls (n = 4); and at 12-15 weeks of age (referred to as “6 weeks”) from *MM2-EGFP* mice (n = 4) and their WT littermate controls (n = 4). Total RNA was isolated from mouse hypothalami using Qiazol lysis reagent followed by purification with the RNeasy Mini kit (Qiagen) according to manufacturer protocol. Genomic DNA contamination was removed with the DNA-free kit (Ambion) and remaining DNA-free RNA was tested for purity using PCR for genomic locus. Total RNA was tested on the 2100 Bioanalyzer (Agilent Technologies) to ensure a RIN quality higher than 9, and quantified using Nanodrop. Equal amounts of total RNA were taken forward for library preparation and ERCC RNA Spike-in control mix (Ambion) was added according to the manufacturer’s guide. Ribosomal RNA was depleted using the RiboErase module (Roche) followed by library preparation using KAPA RNA HyperPrep Kit (Roche). The libraries were sequenced as 50 bp pair-end reads using Nova-seq Illumina platform.

#### Phenotypic analysis

Consistent with previous studies (Brown et al., 2016)(Tillotson et al., 2017), mice were backcrossed for four generations onto C57BL/6J before undergoing phenotypic characterization. Two separate cohorts, each consisting of 10 *MM2-EGFP* animals and 10 wild-type littermates, were produced. One cohort was scored weekly from 4 weeks of age (excluding week 31) until each mutant reached its humane end-point (wild-type controls until 52 weeks) using a system developed by (Guy et al., 2007). Two wild-type controls were culled due to injuries, aged 38 and 45 weeks (censored on survival plot). Mice were scored in six categories: spontaneous activity, gait, hind-limb clasping, tremor, abnormal breathing and general appearance. Mice received a score between 0 and 2 for each category, where 0 = as wild-type, 1 = present, and 2 = severe. Intermediate scores of 0.5 and 1.5 were also used in all categories except hind-limb clasping (Cheval et al., 2012). The scores in each category were added together to give the aggregate symptomatic score for each animal. The mean scores ± S.E.M. for all animals were plotted over time. Animals were also weighed during scoring sessions. Animals were culled if they lost more than 20% of their maximum body weight. Survival was graphed using Kaplan–Meier plots. Previously published (Brown et al., 2016) data for *Mecp2-null*, *T158M-EGFP, R306C-EGFP* and *R133C-EGFP* (all backcrossed onto C57BL/6J) were used for comparison. Scoring: *null* (n = 12), *T158M-EGFP* (n = 11), *R306C-EGFP* (n = 11) *R133C-EGFP* (n = 10); survival: *null* (n = 24), *T158M-EGFP* (n = 11), *R306C-EGFP* (n = 11), *R133C-EGFP* (n = 10); body weight: *null* (n = 20), *T158M-EGFP* (n = 15), *R306C-EGFP* (n = 11), *R133C-EGFP* (n = 10). Scoring data for the *MM2-EGFP*, *R306C-EGFP* and *R133C-EGFP* cohorts was plotted as heatmaps shaded according to the mean score for each category per week (aligned by age and time before death). Two R133C animals survived to one year of age so were excluded from the scoring analysis for the five weeks before death. Some animals reached their humane point >4 days after they were last scored so were included in weeks −5 to −1 only; for 0 weeks prior to death: *MM2-EGFP* (n = 7), *R306C-EGFP* (n = 8), *R133C-EGFP* (n = 8).

The second backcrossed cohorts underwent behavioural analysis at 10-11 weeks of age, consistent with a previous study (Brown et al., 2016). Tests were performed over a two week period: elevated plus maze on day 1, open field test on day 2, hanging wire on day 3 and accelerating rotarod on days 6–9 (1 day of training followed by 3 days of trials). The elevated plus maze is a cross-shaped maze 65 cm above the floor with two open arms (20 × 8 cm), two closed arms (20 × 8 cm, with 25 cm high walls) and a central area (8 × 8 cm). It is set up in a dimly lit room with uniform lighting between each of the closed arms and each of the open arms. Animals were placed in the central area and left to explore for 15 min (during which the experimenter left the room). Mice were tracked using ANYmaze software (Stoelting) and the time in each area was determined. Mice were deemed to be in a particular area of the maze if ≥70% of their body was inside this region. The open field apparatus used consisted of a square arena measuring 50 by 50 cm, which was evenly lit (in a dimly lit room) and littered with fresh wood chippings. The animals were placed in the centre of the area and left to explore for 20 min each (during which the experimenter left the room). Mice were tracked using ANYmaze software (Stoelting). Activity was assessed by total distance travelled. The hanging wire apparatus consists of a 1.5 mm diameter horizontal wire, 35 cm above the bench. Animals were placed on the wire with their forepaws by the experimenter, and the time taken to bring a hind paw to the wire was recorded. Animals were given a maximum of 30 s to complete this task. Animals that took longer or fell off the wire were given the maximum score of 30 s. The test was performed three times for each animal (with an inter-test interval of 30 min) and the mean of the three tests was calculated. The accelerating rotarod consisted of a 3 cm diameter rod that can rotate between 4 and 40 revolutions per minute (rpm). On the first day, the animals were accustomed to the apparatus in a short training session where they must stay on the track for 30 s at its lowest speed (4 rpm). On the subsequent three days, each animal underwent four trials (with an inter-trial interval of 75 min) where the speed is slowly increased from 4 rpm to 40 rpm over 5 min. The time taken to fall (latency) was recorded and the average time for each day is calculated.

All analysis was performed blind to genotype. Animals were randomly assigned to the two backcrossed cohorts and the order in which they were analysed was randomized. Cohorts of this size have been used to successful detect RTT-like symptoms in mice carrying patient mutations including the milder mutation, *R133C* (Brown et al., 2016).

### QUANTIFICATION AND STATISTICAL ANALYSIS

Information on sample size, definition of centre and dispersion, the statistical tests used and the resulting *P* values are included in the figure legends. All statistical analysis was performed using Prism 8 Software. Significance was defined as *P* < 0.05.

#### Evolutionary conservation among MBD proteins

The primary protein sequences of the methyl-CpG binding domains (MBDs) of human and mouse MeCP2, MBD1, MBD2 and MBD4 were aligned using ClustalWS on Jalview 2.8.2.

#### BLI quantification

The binding sensorgrams were analysed using the Octet data analysis software 9.0 (ForteBio, Pall). Experimental data were fitted using a 2:1 model with global fitting (Rmax unlinked by sensors). All experiments were repeated on three independent protein batches and data represents the mean of high-confidence K_D_ (fit R^2^>0.9 and narrow CI). K_D_ values were compared using t-tests (unpaired, two-tailed.

#### EMSA quantification

Percentage shift of the probe in electromobility shift assays (EMSAs) were quantified using Image J (NIH) from three technical replicates.

#### Quantification of TBL1X recruitment

The number of co-transfected cells with TBL1X–mCherry recruitment to heterochromatic foci was determined for each MeCP2 construct. Three independent transfections were performed, and a total of 156, 150 and 151 cells were counted that expressed WT-EGFP, R306C –EGFP and MM2-EGFP, respectively. This analysis was performed blind. The total proportion of cells with TBL1X–mCherry recruitment by each mutant MeCP2 protein was compared with WT using Fisher’s exact tests.

#### FRAP analysis

FRAP analysis was performed on NIH-3T3 cells transiently transfected with EGFP-tagged WT (n = 27) and MM2 (n = 28), over independent three transfections. Images were analysed using a custom plugin (DOI:10.5281/Zenodo.2654602) written for ImageJ (NIH). Fluorescence values were normalised using the following equation: ((Mean Cell Control preBleach-Background(t))/(Control Cell Bleach(t)-Background(t))) * ((FrapSpot(t)-Background(t))/(Mean FrapSpot preBleach-Background(t))). Half-lives were calculated for each cell using the following equation: t_1/2_[S] = t((F’-F0)*0.5 + F0)-t(F0), where F’ = fluorescence maximum at the end of the measurement (mean of the last 10 measurements) and F0 = the fluorescence minimum at t0 (first image after the bleach); logarithmic regression lines were used (calculated in Microsoft Excel, between 0 and 150 s as the curves fitted a logarithmic trend in this portion). The half-lives reported are the mean ± S.E.M. calculated from the replicate cells. The half-lives of WT-EGFP and MM2-EGFP where compared using a Mann-Whitney test.

#### Quantification of Flow Cytometry

Protein levels in neuronal nuclei were quantified by EGFP fluorescence by Flow cytometry (BD LSRFortessa SORP using FACSDIVA version 8.0.1 software) to obtain the mean EGFP fluorescence for the total nuclei (n = 50,000 per sample) and the high-NeuN (neuronal) subpopulation (n > 8,000 per sample). Three biological replicates of each genotype were analysed. The protein levels in MM2-EGFP mice were compared with WT-EGFP controls using t-tests (unpaired, two-tailed).

#### Quantification of ChIP-qPCR

To quantify ChIP-qPCR, the efficiency of each primer pair was determined from the standard curve and this value was used to calculate IP/input (%). Enrichment in WT-WGFP and MM2-EGFP samples (n=3 biological replicates per genotype) were compared using t-tests (unpaired, two-tailed).

#### Quantification of genomic cytosine methylation

To calculate the percentage methylation in the Whole genome bisulphite sequencing data (GEO: GSE84533) (Lagger et al., 2017), a threshold of ≥5-fold coverage was used. The proportion of methylation for each site was determined based on the reads covering the site for each motif, and the sum of these values was divided by the number of time that motif occurred. To determine the number of sites with different levels of methylation (100%, 75-99%, 50-74%, 25-49%, 1-24% and 0%), a threshold of ≥5-fold coverage was used. To calculate the approximate number of sites with different levels of methylation in the whole genome, these values were multiplied by total number of motifs in the genome/number of motifs with ≥5-fold coverage.

#### Analysis of ATAC-seq footprinting

Nextera adapter sequence removal was performed on 75 bp paired-end reads using Trimmomatic version 0.32 with the following parameters: ILLUMINACLIP:nextera.fa:2:40:15 MINLEN:25. Surviving reads were then mapped to the mouse mm9 reference genome with bwa mem version 0.7.5a-r405 using the −M parameter. Alignments were filtered based on mapping quality (<20) so that only uniquely mapped reads were taken forward, furthermore, reads aligning to mitochondrial DNA or Encode mm10 blacklisted regions were removed from further analysis. Read alignments were converted into normalised coverage files for visualisation and downstream quantification using deepTools bamCoverage version 3.1.3 with the following parameters: -bs 1 --normalizeUsing BPM --skipNAs -e. The positions of Tn5 transposase cutsites were inferred from the 5’ end of aligned sequencing reads and converted into bigWig format for further analysis.

Whole genome bisulphite sequencing data (GEO: GSE84533)(Lagger et al., 2017) was employed to visualise a footprint of MeCP2 binding across methylated and unmethylated cytosines in several sequence contexts. For mCG, 1 million randomly selected sites with 100% methylation (≥5-fold coverage between both strands) were used. For CG, sites with 0% methylation (≥10-fold coverage between both strands) were used (n = 935,582). For mCAC, sites with ≥75% methylation (≥5-fold coverage) were used (n = 95,017). To increase the number of sites used, the thresholds were lowered to ≥4-fold coverage and ≥50% methylation for other forms for mCH: mCAT n = 167,159; mCAG n = 172,250; mCAA n = 72,688; mCT n = 263,578; mCC n = 7,643. For all unmethylated CH motifs, 1 million randomly selected sites with 0% methylation (≥10-fold coverage) were used. The distribution of ATAC-seq cut sites around methylated and unmethylated cytosines was computed in R using the seqPlots package. The region 300 bases upstream and downstream of each cytosine was divided into single base bins and for each bin the total number of cut sites was summed across all cytosines. These values were plotted to show the number of cut-sites in the 600 bases surrounding cytosines of different methylation context. The values were normalised to the flanking regions by calculating the ratio of each bin to the mean of all bins within the outer 100 bases of the plot. For each sample the mean values of each bin were computed across all biological replicates before plotting. To determine footprints of MeCP2-dependent or MM2-EGFP-dependent chromatin inaccessibility, ATAC-seq profiles over each motif were divided by the equivalent KO profile. Graphs show ln(WT/KO) or ln(*MM2-EGFP*/KO).

#### Analysis of RNA-seq datasets

Raw sequencing reads are quality and adapter trimmed using Trimmomatic v0.33 (Bolger et al., 2014). Quality trimmed reads are aligned to mm10 genome assembly and M19 transcriptome assembly using STAR v2.4.2a (Dobin et al., 2013) and read counts spanning exons are calculated using featureCounts v1.6.4 (Liao et al., 2014). Differential gene expression analysis is performed using DESeq2 v1.24.0 (Love et al., 2014) and significance threshold for a gene is set at p-adjusted value < 0.05.

Whole genome bisulphite sequencing data (GEO: GSE84533)(Lagger et al., 2017) was used to determine the number of mCG and mCAC binding sites per gene. Since a single molecule of an MBD protein binds a symmetrically methylated mCG/mCG site, this number was divided by two. Comparisons of gene expression changes to number of methylated binding sites used all genes that were detected in all datasets (n = 14,391). Gene expression changes were determined by comparing the following pairs of datasets: KO/WT (littermate controls, at 6 weeks on C57BL/6J;CBA/CA), *MM2-EGFP*/WT (littermate controls, at 6 weeks on C57BL/6J), *MM2-EGFP*/WT (littermate controls, at 12 weeks on C57BL/6J). Genes were binned by the number of mCG or mCAC binding sites, each bin contained 500 genes (the first and last bins were excluded due to the large range of the binned variable). Slopes of linear regression (with 95% confidence interval) were determined and plotted for each bin. The ratio of mCG/mCAC binding sites was calculated for genes in each of the mCAC bins and the ratio of mCAC/mCG binding sites was calculated for genes in each of the mCG bins.

Genes significantly dysregulated in mutants compared to age-matched wild-type littermate controls were identified in each dataset (KO/WT, 6 weeks; *MM2-EGFP*/WT 6 weeks; and *MM2-EGFP*/WT, 12 weeks) using an adjusted p-value (padj) of <0.05. Comparisons between the shared differentially expressed genes and those dysregulated in KO or MM2-EGFP only were performed separately on the age-matched and symptom-matched datasets. This analysis compared: the total number of mCG + mCAC binding sites per gene, the mCAC/mCG ratios per gene and expression changes (relative to wild-type littermate controls). Shared genes were statistically compared to those affected in KO or MM2-EGFP only using Mann-Whitney tests. Expression changes of the neurological disease-associated genes from previously published RNA-seq datasets were added for comparison: KO/WT hypothalamus 6 weeks (GEO: GSE66870)(Chen et al., 2015) and KO/WT cortex 8 weeks (GEO: GSE128178)(Boxer et al., 2019).

#### Disease Ontology Analysis

Dysregulated gene identifiers are converted to their *H. sapiens* orthologues and disease ontology enrichment analysis is performed using clusterProfiler (Guangchuang Yu, Li-Gen Wang, Yanyan Han, Qing-Yu He. clusterProfiler: an R package for comparing biological themes among gene clusters. OMICS: A Journal of Integrative Biology. 2012, 16(5):284-287.) Over representation analysis (ORA) is used to determine whether any known disease ontologies are over-represented in the list of dysregulated genes. Hypergeometric test is used to estimate the significance of the disease ontologies and p-values are adjusted to account for Type I errors using Bonferroni correction. Type I errors are also accounted using a separate independent method called Benjamini/Hochberg correction to calculate false-discovery rates (FDR).

#### Statistical analysis of behavioural data

Growth curves were compared using Mixed-effects model (REML) analysis for weeks 4-48 (excluding week 31). Survival curves were compared using a Mantel–Cox test. For behavioural analysis, when all data fitted a normal distribution (open field), genotypes were compared using t-tests (unpaired, two-tailed). For non-parametric datasets (elevated plus maze, hanging wire and accelerating rotarod latency to fall), genotypes were compared using Kolmogorov–Smirnov tests. Change in performance over time in the accelerating rotarod test was determined for each genotype using Friedman tests.

### DATA AND SOFTWARE AVAILABILITY

FRAP macro script: DOI:10.5281/Zenodo.2654602

ATAC-seq and RNA-seq: GEO GSE152801

### KEY RESOURCES TABLE

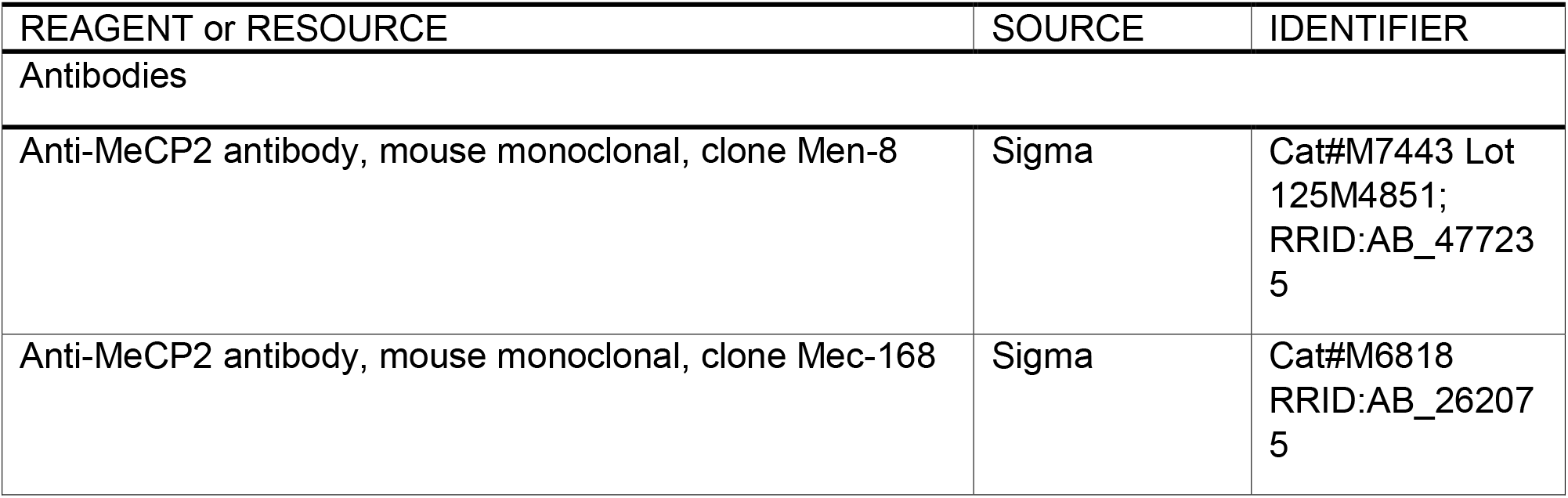
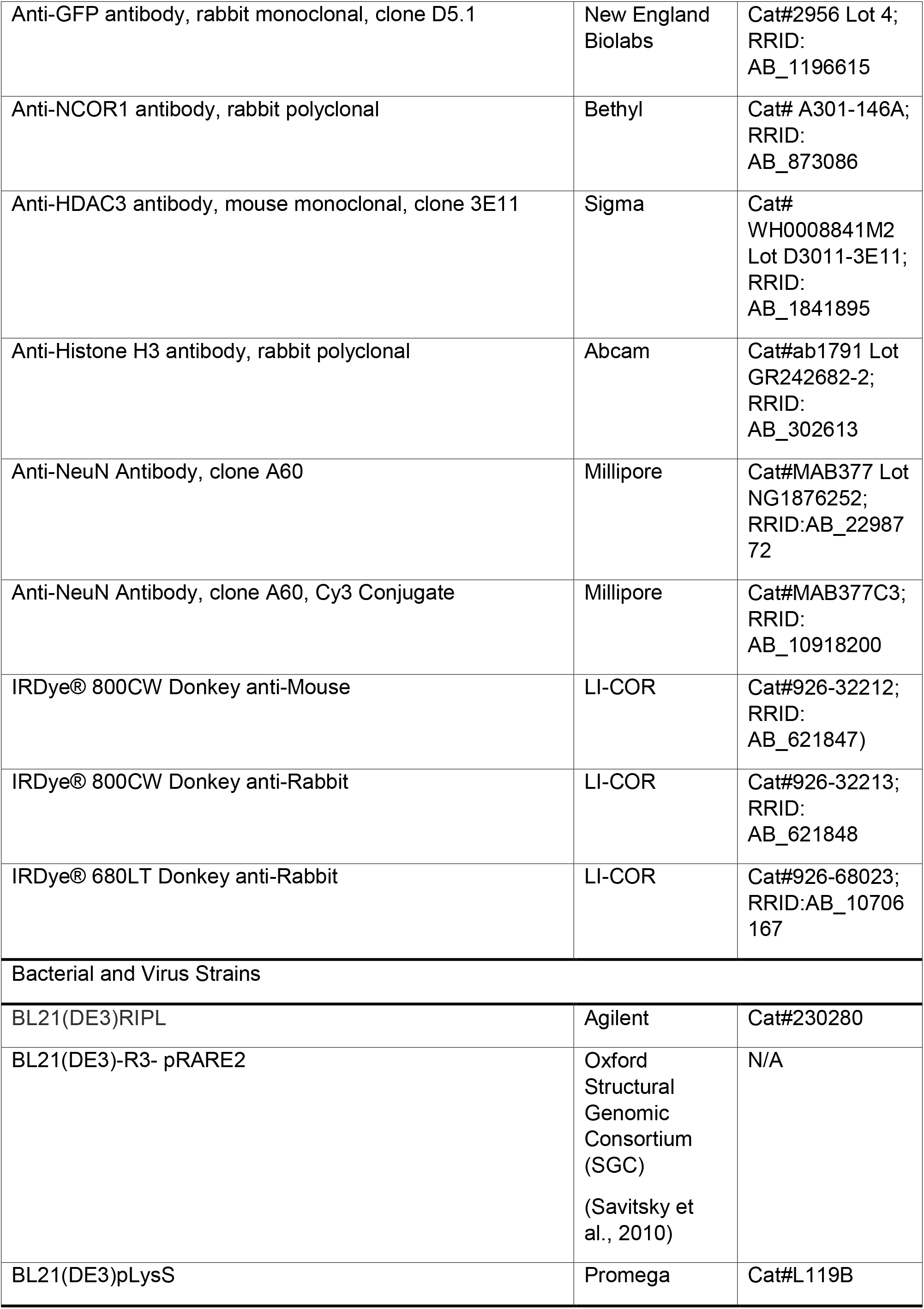
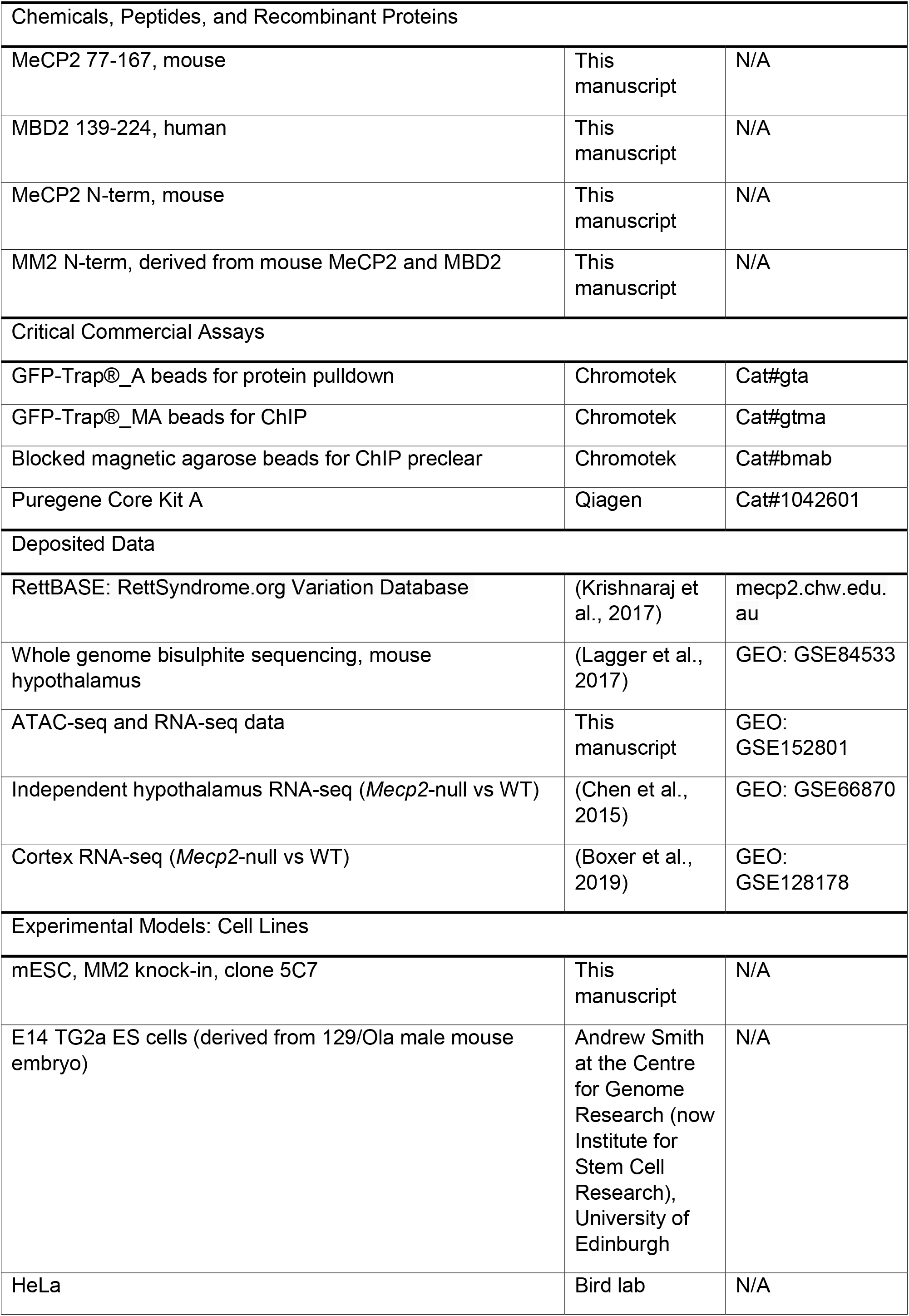
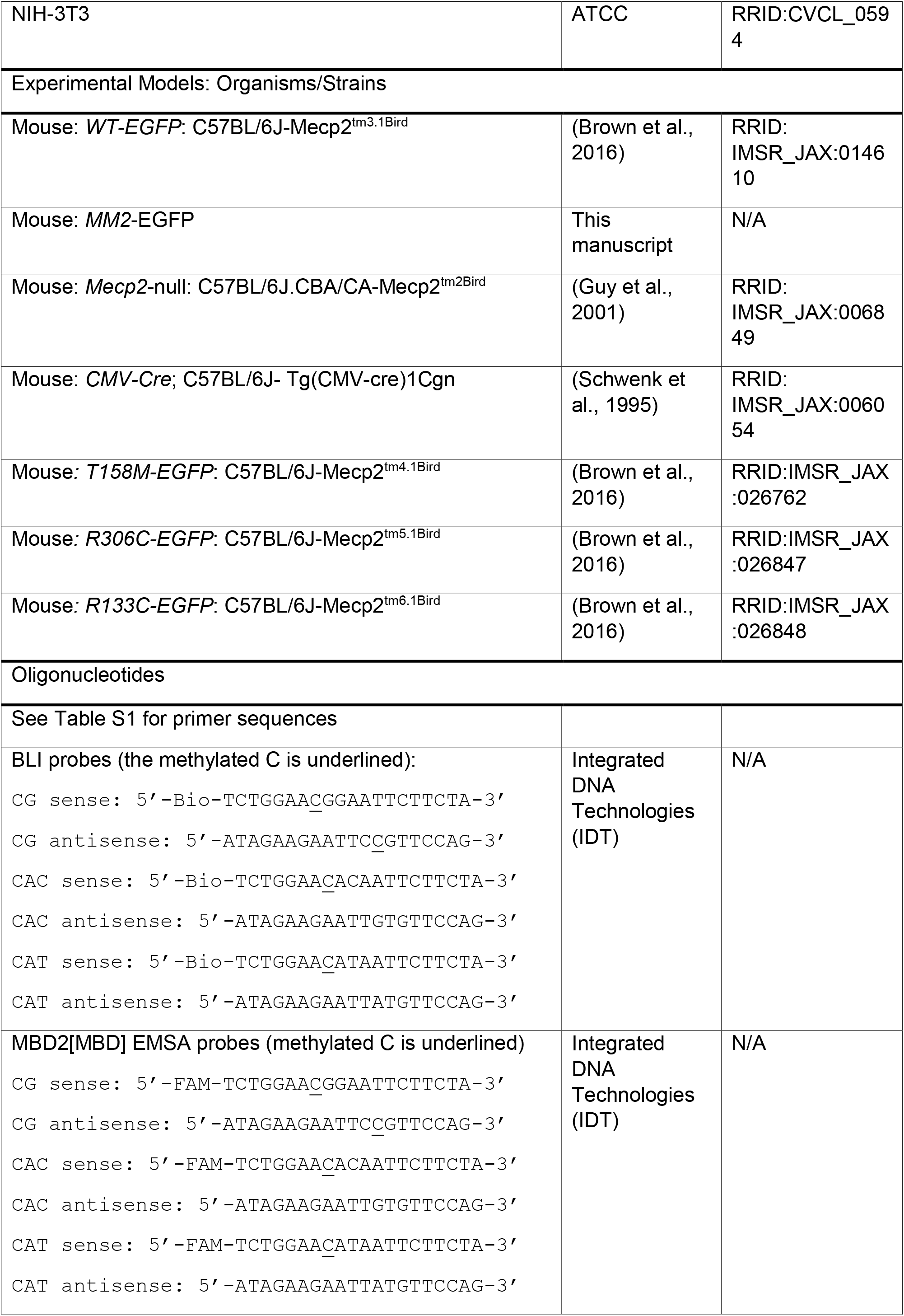
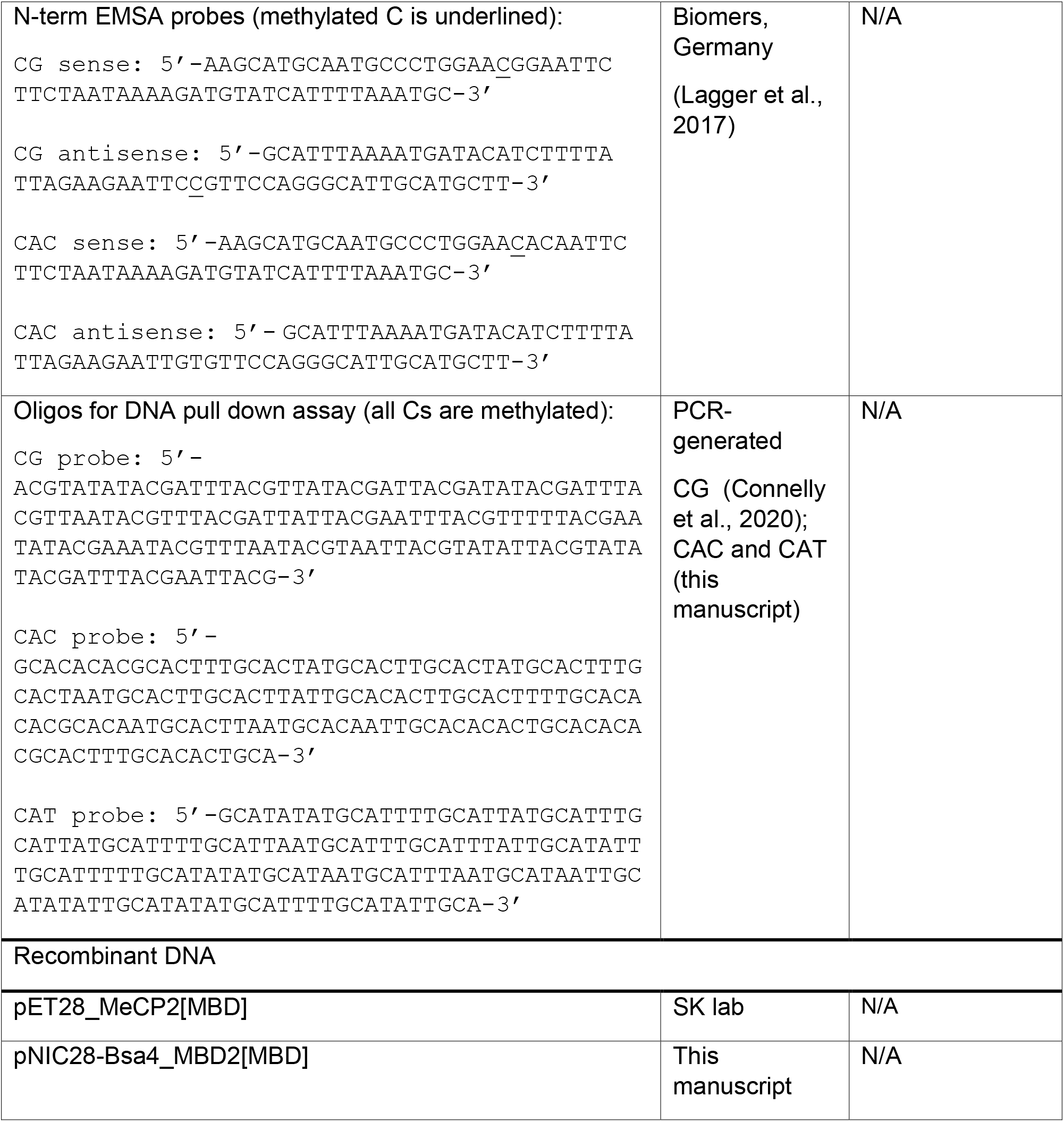
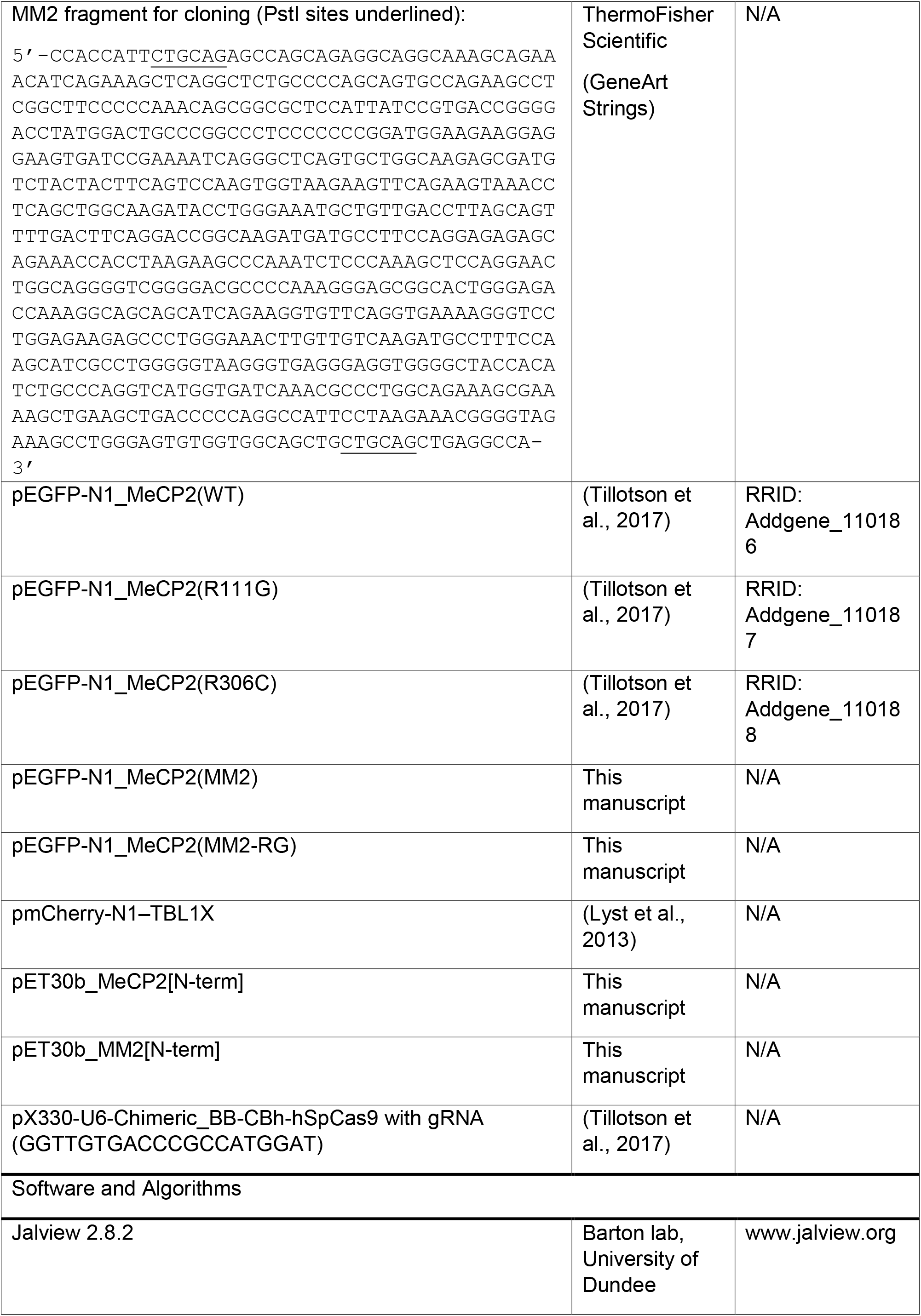
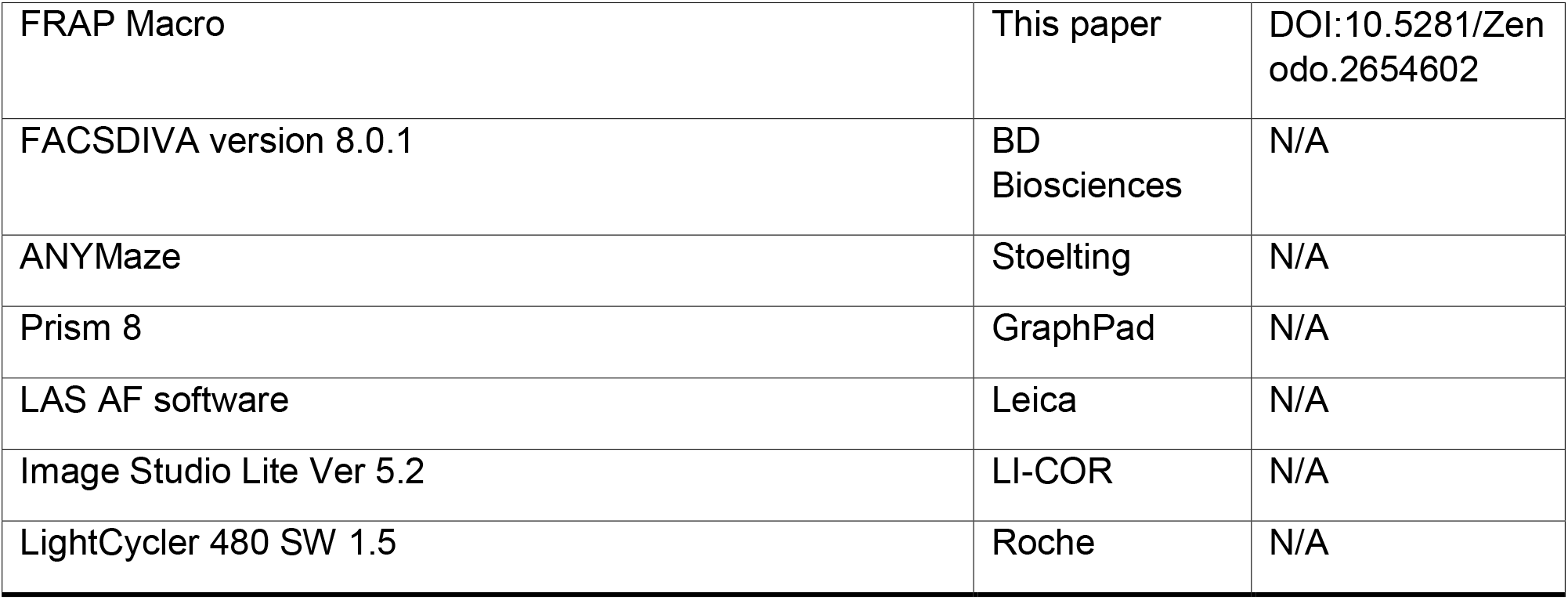

**Table S2:**
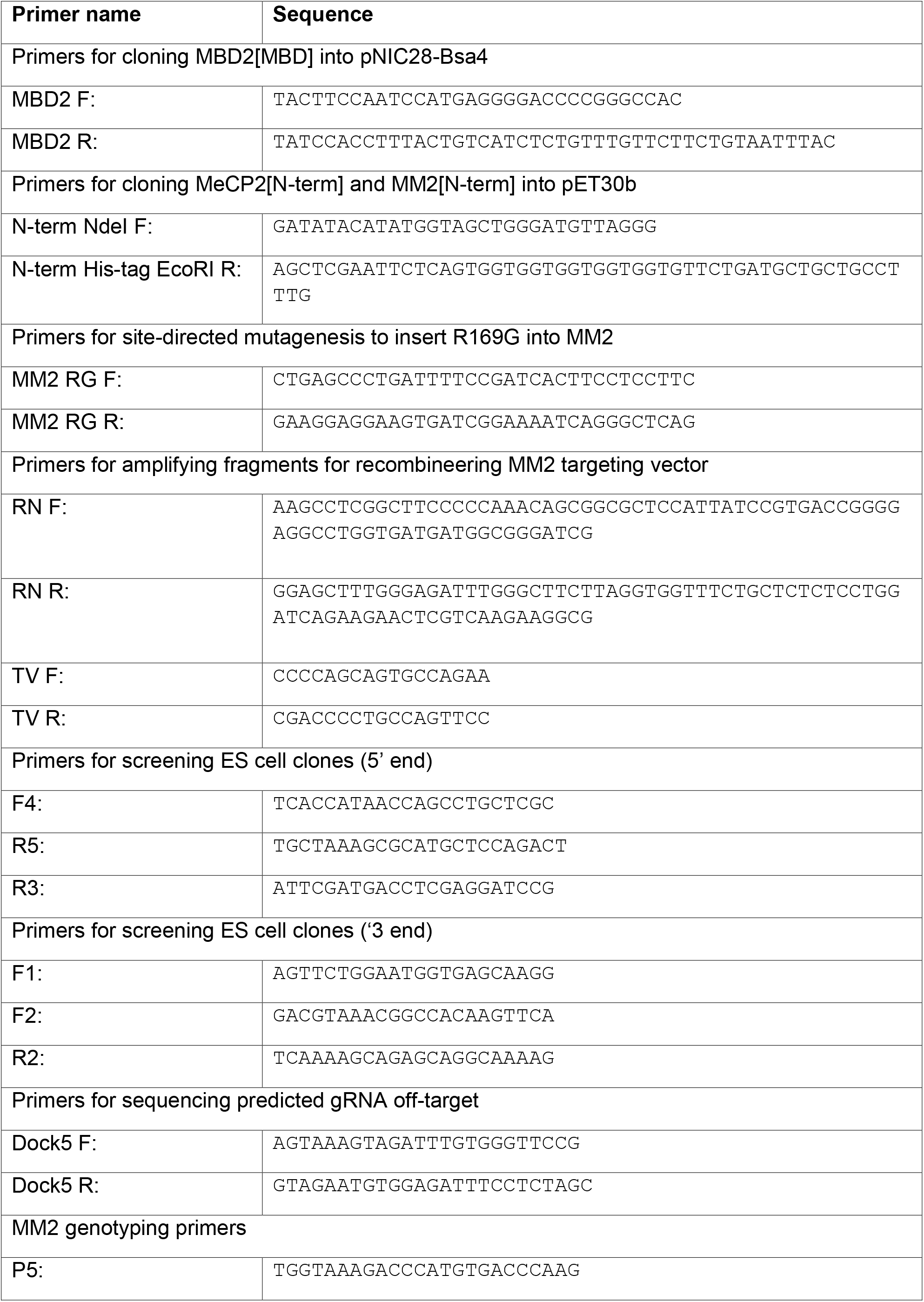
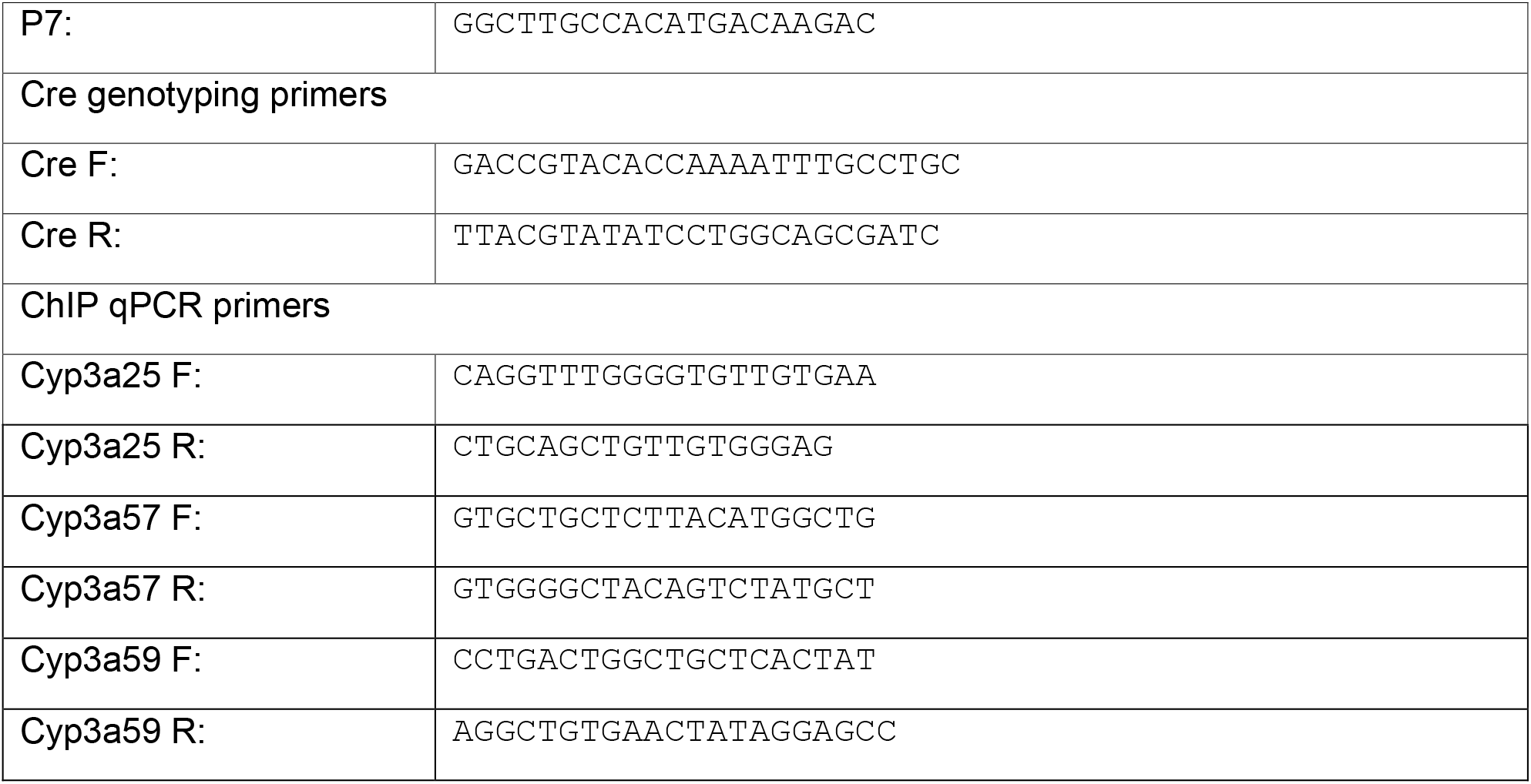
Primer sequences.

